# Microbial Respiration in Contrasting Ocean Provinces via High-Frequency Optical Assays

**DOI:** 10.1101/2023.07.20.549894

**Authors:** Melanie R. Cohn, Brandon Stephens, Meredith G. Meyer, Garrett Sharpe, Alexandria K. Niebergall, Jason R. Graff, Nicolas Cassar, Adrian Marchetti, Craig A. Carlson, Scott Gifford

## Abstract

Microbial respiration plays a pivotal role in the marine carbon cycle, influencing the fraction of fixed carbon that undergoes remineralization versus export to depth. Despite its importance, methodological constraints have led to an inadequate understanding of this process, especially in low-activity oligotrophic and mesopelagic regions. Here, we quantify respiration rates as low as 0.2 *µ*mol O2 L^-1^ d^-1^ in contrasting ocean productivity provinces using optical oxygen sensors to identify size-fractionated respiration trends. At the low productivity North Pacific Ocean Station Papa, surface microbial respiration was relatively stable at 1.2 *µ*mol O2 L^-1^ d^-1^. Below the surface there was a decoupling of respiration and bacterial production potentially driven by phytodetritus remineralization. Size-fractionated rates showed cells <5 *µ*m contributed the most to Pacific respiration. At the North Atlantic Porcupine Abyssal Plain, the optode measurement frequency was drastically increased. Surface microbial respiration was higher (1.7 *µ*mol O2 L^-1^ d^-1^) and decreased by 3-fold below the euphotic zone. The Atlantic filtered fraction contributions to total respiration shifted with the phytoplankton bloom evolution. The high resolution optode method used in the Atlantic is consistent with coinciding *in vivo* 2-para-(iodophenyl)-3(nitrophenyl)-5(phenyl) tetrazolium chloride respiratory stain measurements and historical site estimates. We estimate that 58% of gross primary production was respired at the Pacific site and 34% at the Atlantic site, demonstrating that the Atlantic had a higher carbon export potential. This study highlights the dynamic nature of respiration across vertical, temporal, and size-fractionated factors, emphasizing the need for sensitive, high-throughput techniques to better understand ocean ecosystem metabolism.

## INTRODUCTION

Aerobic respiration is a crucial process in which organic carbon is oxidized to fuel metabolism, transforming the carbon pool and generating carbon dioxide (CO2). In the ocean, up to half of all phytoplankton-derived organic matter is respired by heterotrophic bacteria (Williams 1984), which reduces carbon export potential (Azam et al. 1983; Moran et al. 2022). However, predicting microbial respiration rates is challenging due to complicating factors such as environmental conditions, community composition, and organic substrate types and availability, making it difficult to understand how shifts in the ecosystem alter respiration rates (del Giorgio et al. 1997; Robinson et al. 2002). Furthermore, respiration rates in the ocean are poorly constrained due to under-sampling and methodological limitations, reducing the accuracy of biogeochemical models (Pomeroy et al. 1995; del Giorgio and Duarte 2002; Robinson 2019). The uncertainties are great enough such that there is debate whether the oligotrophic oceans are net autotrophic or net heterotrophic (Ducklow and Doney 2013 and references therein), hindering our ability to predict the impact of climatological stressors on ocean ecosystems.

Respiration rates are often determined via incubation assays in which oxygen (O2) drawdown is measured over time. Accurate measurements are hard to obtain given the relatively small change in O2 (<1 *µ*mol L^-1^) against the large background concentration in seawater (*c.a.* 200 - 300 *µ*mol L^-1^). As a result, most methods rely on gathering a few highly precise O2 measurements at the cost of sample throughput. Traditional Winkler titrations obtain accurate and precise dissolved O2 measurements, but are time consuming, destructive to the sample and resource intensive. Respiratory stains such as *in vivo* 2-para-(iodophenyl)-3(nitrophenyl)-5(phenyl) tetrazolium chloride (INT) measure activity of the electron transport system, yielding highly precise and sensitive measures of respiration, but face similar challenges as Winkler titrations in terms of time and wet lab resources (Griffith 1988).

Optical and electrochemical approaches are increasingly common methods to determine dissolved O2 concentrations for respiration and net community production in marine environments. Previously, these assays were often unable to resolve rates <1 *µ*mol O2 L^-1^ d^-1^ without concentration of environmental samples (Pomeroy and Johannes 1966; Griffith 1988; del Giorgio and Williams 2005). Edwards et al. (2011) were able to reliably measure rates below 1 *µ*mol O2 L^-1^ d^-1^ using O2 optodes and incubation assays. Wikner et al. (2015) further expanded on this, showing optode detection limits three times lower, comparable to Winkler titrations. The sensitivity and ease of use of foil-membrane optodes are promising, as they are capable of rapid, frequent point measurements (on the order of seconds). Rather than taking relatively few point measurements hours apart, hundreds of measurements can be made through high-temporal resolution assays, with the potential to resolve pelagic respiration rates under shorter incubations (Chen et al. 2023). Through improved sample throughput and coverage, optical methods have the potential to provide needed insight into the critical role of microbial respiration in global, oceanic carbon cycling (Edwards et al. 2011; Martínez-García and Karl 2015; Robinson 2019).

In this study, we determined microbial respiration rates via O2 drawdown assays using high-frequency O2 measurements from continuously-mounted optodes in two oceanic provinces: the low productivity and tightly coupled N. Pacific Ocean Station Papa (Wong et al. 1995; Boyd and Harrison 1999; Meyer et al. 2022) and the high productivity N. Atlantic Porcupine Abyssal Plain (PAP) (Fasham et al. 1999; Frigstad et al. 2015). This work is part of the EXport Processes in the Ocean from RemoTe Sensing (EXPORTS) program to determine a carbon cycle budget across ocean sites at opposing ends of the productivity-export spectrum (Siegel et al. 2016, 2021) and capturing the N. Atlantic spring bloom at heightened activity compared to other times of the year (Meyer et al. 2023). Through the EXPORTS field campaigns in both provinces, we explore the role of microbial metabolism in oceanic carbon cycling and export potential. Here, we implemented optode-derived respiration measurement to constrain respiration rates at low and high productivity endmember systems within and below the euphotic zone, for different size fractions, and in response to physical and biological changes in environmental states. We demonstrate the power of optodes and incubation assays to resolve respiration rates in pelagic systems with reduced ship-board effort. Our results show sample size-fractionation differences in microbial community contributions to total respiration between surface and sub-euphotic depths, as well as across distinct oceanic provinces. Microbial respiration is relatively stable in the low-activity N. Pacific with over half of gross primary production (GPP) respired. It is highly dynamic on daily timescales in the N. Atlantic, correlating with biomass and particle trends in this greater carbon export system.

## METHODS

### Study site and sampling

Samples were collected in the N. Pacific at Ocean Station Papa (Station P; 50° N and 145° W) from August 8 to September 3, 2018 aboard the R/V *Roger Revelle* (RR1813) and in the N. Atlantic at the Porcupine Abyssal Plain (49° N, 15° W) from May 5 to 31, 2021 aboard the RRS *James Cook* (JC214) while following Lagrangian floats (Siegel et al. 2021; Johnson et al. 2023) (Fig. 1). Sample water was collected from 10-L Go-Flow (N. Pacific) or 20-L Niskin (N. Atlantic) bottles attached to a CTD rosette directly into 1-L polycarbonate bottles (acid-washed with 10% HCl). CTD casts were deployed between 2 to 5 am GMT in the N. Atlantic and variable times in the N. Pacific. At minimum for each CTD sampling, a “surface” depth (<10 m in the N. Pacific; 5 m in the N. Atlantic) was collected. In the Pacific, depth profiles to 200 m were opportunistically taken, and in the Atlantic a “sub-euphotic” depth (collection depth of 75 m or 95 m; euphotic zone range of 1% irradiance from 37 to 59 m) near the base of the mixed layer (range 15 m to 197 m, 0.03 kg m^-3^) was collected (Meyer et al. 2023). Additional physical context methodology is referenced in the ancillary data below.

**Figure 1.**
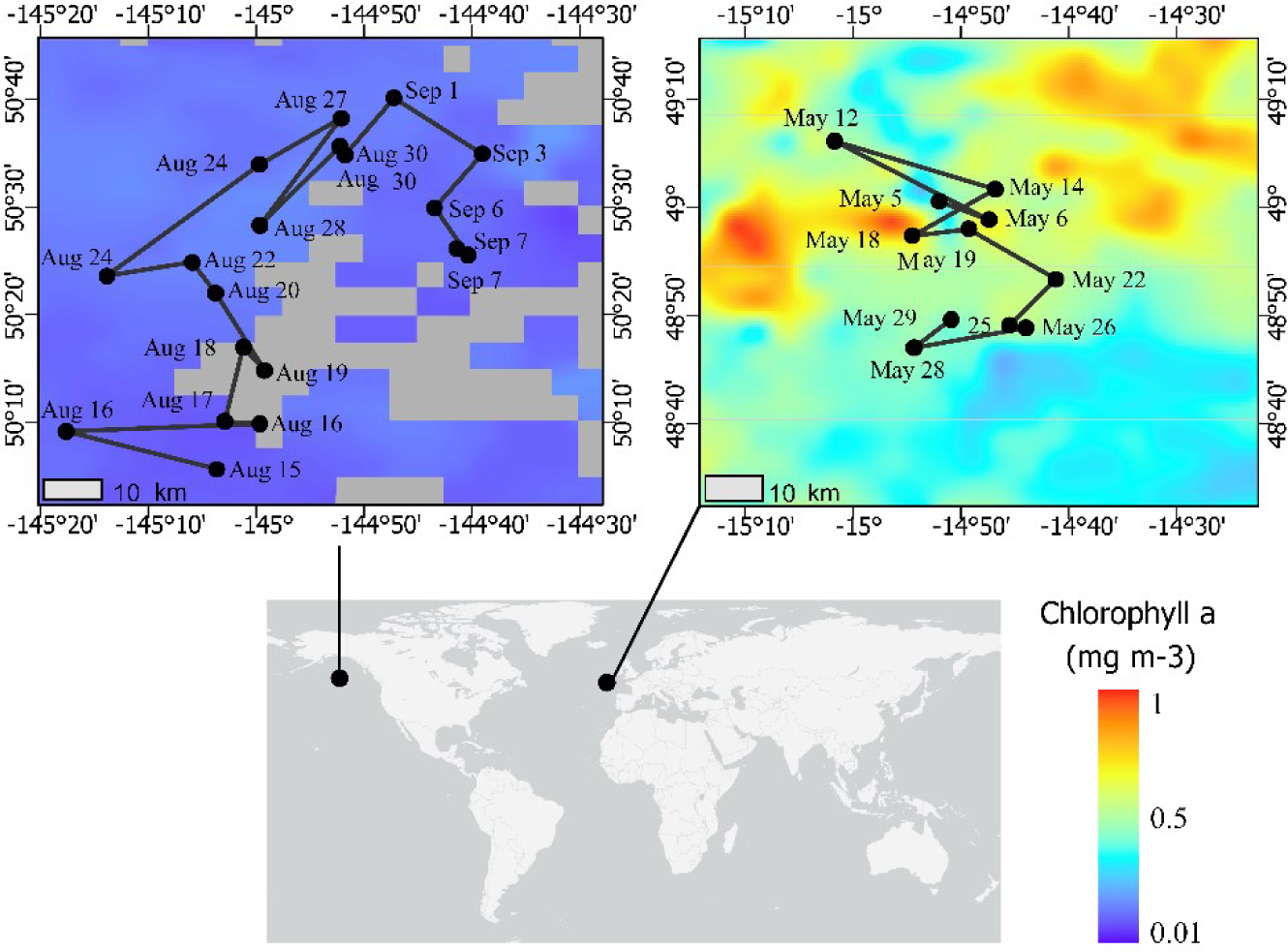
Map of EXPORTS field campaigns for August 2018 at Ocean Station Papa and May 2021 at the Porcupine Abyssal Plain. Sampling dates are labeled on the Lagrangian (Pacific) and quasi-Lagrangian (Atlantic) cruise tracks. The color scale depicts the monthly average of chlorophyll *a* at each site during the occupation period (NASA MODIS-AQUA, https://oceancolor.gsfc.nasa.gov/l3/).

### Seawater sample handling and preparation

Whole or size-fractionated seawater incubations were performed in 60-ml glass biological oxygen demand (BOD) bottles with O2 sensor spots (SP-PSt-3-NAU, PreSens) mounted to the interior of the bottle wall. The BOD bottles were washed with 5% HCl for 3-6 h and triple rinsed with 18.2 MΩ Milli-Q water before use. For size-fractionation, seawater samples were poured from the collection bottle into acid-washed polycarbonate filter towers. The tower and filter were rinsed with 100 mL of seawater sample, the rinsate was discarded, then 300 mL of sample was filtered under gentle (>-5 in Hg) vacuum pressure. Size-fractions in the N. Pacific included unfiltered and <5 *µ*m (polycarbonate, Millipore Sigma). The N. Atlantic had these fractions plus <3 *µ*m and <1.2 *µ*m fractions (polycarbonate, Millipore Sigma). The BOD bottles were then filled with either unfiltered or size-fractionated sea water samples by first triple rinsing with *c.a.* 20 mL of sample water poured directly from the collection bottle (unfiltered) or filter tower (size-fractionated) and filled to the brim. Bubbles were removed from the BOD bottle over one minute by gently tapping the side, then the bottle was capped with an air-tight glass stopper.

The BODs were submerged in a dark, temperature-controlled water bath at *c.a. in situ* temperature for incubation. Incubation times ranged from 16 to 72 h based on detection of a robust decrease in O2 concentration (Vandermeulen and Chaves 2022). In the N. Pacific, O2 measurements were made non-invasively through the bottle wall at 4 h intervals by manually holding the optical fiber to the BOD wall at the sensor spot (OXY1-SMA, PreSens). The N. Pacific average incubation time was 38 h with 8 to 10 discrete O2 measurements per assay. Over the 30-day occupation, 214 O2 drawdown assays were conducted at Station Papa with 94 unfiltered and small (<5 *µ*m) size fraction assays in the mixed layer (avg. 29 ± 4.5 m; Siegel et al. 2021). In the N. Atlantic, O2 measurements were acquired continuously every 30 seconds by directly mounting the polymer optical fibers to the BOD bottles over the sensor spot. Measurements were made with OXY4-SMA (PreSens), with an average incubation time of 30 h and 3,000 discrete O2 measurements per assay, totaling 41 unfiltered and 66 filtered assays.

### O2-based respiration rate determinations

O2 measurements were automatically temperature and salinity corrected through the PreSens Measurement Studio2 software based on direct, continuous temperature measurements of the water bath and a defined *in situ* salinity and pressure (35 psu, 1013 hPa). For each incubation, a BOD bottle with sensor spot was included that had been acid washed, rinsed, and filled with Milli-Q water to serve as an additional temperature control and to monitor sensor drift. This control bottle was co-incubated with the samples and had a dedicated mounted optode. Further temperature correction of the samples was performed by subtracting the change in O2 concentration of a Milli-Q control bottle from the sample bottles for each point. The corrected sample O2 values were then plotted against incubation time.

For the N. Pacific dataset in which optode measurements were manually acquired, there were relatively few O2 measurements per experiment (*n*=8 to 20). The O2 drawdown rates were calculated as the Model I regression of the measured points. Apparent O2 outliers (>1 *µ*mol L^-1^ O2 difference from the surrounding points) were identified and removed within each assay (avg. 1.2 points removed). Assays with rates <0.2 *µ*mol O2 L^-1^ d^-1^ were considered below detection (*n*=9) and rates greater than 3 *µ*mol O2 L^-1^ d^-1^ were considered anomalously high for summer at the Pacific site and excluded from analysis (*n*=3).

In the N. Atlantic where the continuously mounted optical fiber cables were used, the first 2.5 h of data were removed to compensate for abiotic acclimation of the PSt3 sensor spot and BOD bottles to the water bath conditions. Plots were visually inspected to trim non-linear components at the beginning or end of the incubation, similar to effects discussed in Wikner et al. (2013). To calculate the rate over *c.a.* 3,000 discrete O2 measurements in a single assay, a rolling mean with a window size of 12.5 min was applied to the trimmed data, and a linear regression was calculated based on the rolling mean points.

### INT-based respiration rate determinations

Respiratory stain (INT) and O2-based incubations were conducted independently on the EXPORTS N. Atlantic expedition and compared. Sample water for INT incubations was collected at midday casts whereas O2 drawdown samples were collected from pre-dawn CTD casts. For each depth, 3 live sample and 2 poisoned (2 w/v formaldehyde) control incubations were conducted. Following methods detailed by Martínez-García et al. (2009), 8 mmol L^-1^ INT was added to a final concentration of 0.2 mmol L^-1^ and incubated in the dark at *in situ* temperature for 2 h. The samples were then filtered under gentle vacuum pressure through stacked, in-line filters (45 mm 3.0 *µ*m and 47 mm 0.2 *µ*m, polycarbonate, Millipore Sigma) to capture size-fractionated cells. The absorbance of the extracted propanol-formazan solution was measured at 485 nm. The INT-based rate is converted to O2 consumption using 3 unique 12-point dilution-based standard curves of freshly prepared Fomazan dye (Fisher Scientific). To compare the O2 drawdown method to the INT methods, rates were selected from both methods where sample collection occurred within 16 h of each other.

### Diel phytoplankton-derived carbon experiment

Water from 5 m at PAP in the N. Atlantic was collected from a CTD cast into 2-L, acid-washed, clear polycarbonate bottles and filled with no headspace on May 29, 2021. The bottles were placed in an on-deck incubator at 40% irradiance to replicate *in situ* phytoplankton growth. Every 6 h for 2 days, one bottle was subsampled and filtered through a rinsed, 1.2-*µ*m filter. Duplicate BOD bottles were filled with the filtrate and placed in the dark respiration water bath at *in situ* temperature for respiration assays.

### Organic carbon contamination monitoring

Care was taken to limit carbon contamination in respiration assays by acid washing and sample water rinsing surfaces as well as using gentle vacuum filtration to reduce cell shear. All respiration assays were monitored for anomalously high rates (>3 *µ*mol L^-1^ d^-1^). On intermittent days, additional sample replicates were prepared such that randomly selected BOD bottles for each depth and size fraction were sampled at the start and end of the incubations for total organic carbon (TOC) or dissolved organic carbon (DOC) analysis as described in Stephens et al. (2020). These respiration incubations were run longer than our standard assays (>2 d, until the change in O2 concentration was at least 2.5 *µ*mol L^-1^ O2) with the goal of allowing for a measurable DOC drawdown to compare to O2 drawdown rates.

### Primary production

Chlorophyll *a*, net and gross primary production were performed in the Pacific as described in Meyer et al. (2022) and in the N. Atlantic as described by Meyer et al. (2023). Gross primary production (6 h ^13^C-uptake incubation) and net primary production (NPP, 24 h ^13^C uptake incubation) were estimated based on samples collected at the surface on the same days as respiration assays in the N. Pacific and at the same CTD casts as respiration assays in the N. Atlantic. The percent of GPP respired was calculated by averaging the daily surface ratios of unfiltered respiration rates to volumetric GPP rates. For this calculation, respiration was converted to carbon units using a respiratory quotient of 1.4 O2 to CO2 (Laws 1991; Stephens et al. 2023).

### Statistical analysis

Linear regressions were performed on each respiration assay as described above using a general linear regression model with the coefficients of determination reported (Fig. S2, S3). For the N. Atlantic, the residual standard error (RSE) was calculated across the multiple size fractions and unfiltered samples. One outlier was identified in the sub-euphotic <5*µ*m fraction using the ROUT method (Q=1%) and removed. Welch’s ANOVA and Dunnett T3 multiple comparisons test (*α*=0.05) were used to identify significantly different (*p*<0.001) RSE pairings across size fractions and depth (Fig. S5, S6). Pearson’s correlation coefficients were calculated on the means and standard deviations of the N. Atlantic biological variables, assuming Gaussian distributions to identify significant (*n*=8 to 18, *p*<0.05) relationships (Fig. S7).

### Ancillary data and methods

Physical parameters were defined by Siegel et al. (2018) in the N. Pacific; and Erikson et al. (2023) and Johnson et al. (2023) in the N. Atlantic. Bacterial abundance was enumerated via flow cytometry as described in Stephens et al. 2020. To evaluate bacterial activity, ^3^H-leucine incorporation rates are used as a proxy for bacterial production (BP) as described by Stephens et al. 2020. Phytoplankton were enumerated via flow cytometry following Graff and Behrenfeld (2018).

## RESULTS AND DISCUSSION

### N. Pacific respiration rates

Station Papa is a high nitrate, low chlorophyl region where primary productivity was low and representative of typical conditions during the sampling (Fig. 1; Boyd and Harrison 1999; Meyer et al. 2022). Fittingly, we observed low total respiration rates in this open ocean environment in late summer 2018. Over the 30-day occupation, 214 O2 drawdown assays were conducted at Station Papa (Fig. 1) with 94 unfiltered and small (<5 µm) size fractions assays in the mixed layer (avg. 29 ± 4.5 m (Siegel et al. 2021)). Over the two-day incubation period, typically 8 to 10 O2 measurements were obtained for each assay by manually holding the optic fiber over the sensor spot. Linear fits of drawdown over time were variable, ranging from strong (*r^2^*>0.8, *n*=25), to moderate (*r^2^*=0.5 to 0.8, *n*=33), and weak (*r^2^*<0.5, *n*=12) (Fig. 2 b-d, Fig. S1). Surface respiration rates averaged 1.2 *µ*mol O2 L^-1^ d^-1^ for both the unfiltered (range 0.34 to 2.25 *µ*mol L^-1^ O2 d^-1^) and small fractions (range 0.55 to 2.41 *µ*mol L^-1^ O2 d^-1^) and were relatively consistent within error (Fig. 2a). These rates are similar to previously measured mean 1.56 *µ*mol O2 L^-1^ d^-1^ total respiration rate derived from Winkler titrations in summer 1997 at Station Papa (Sherry et al. 1999).

**Figure 2.**
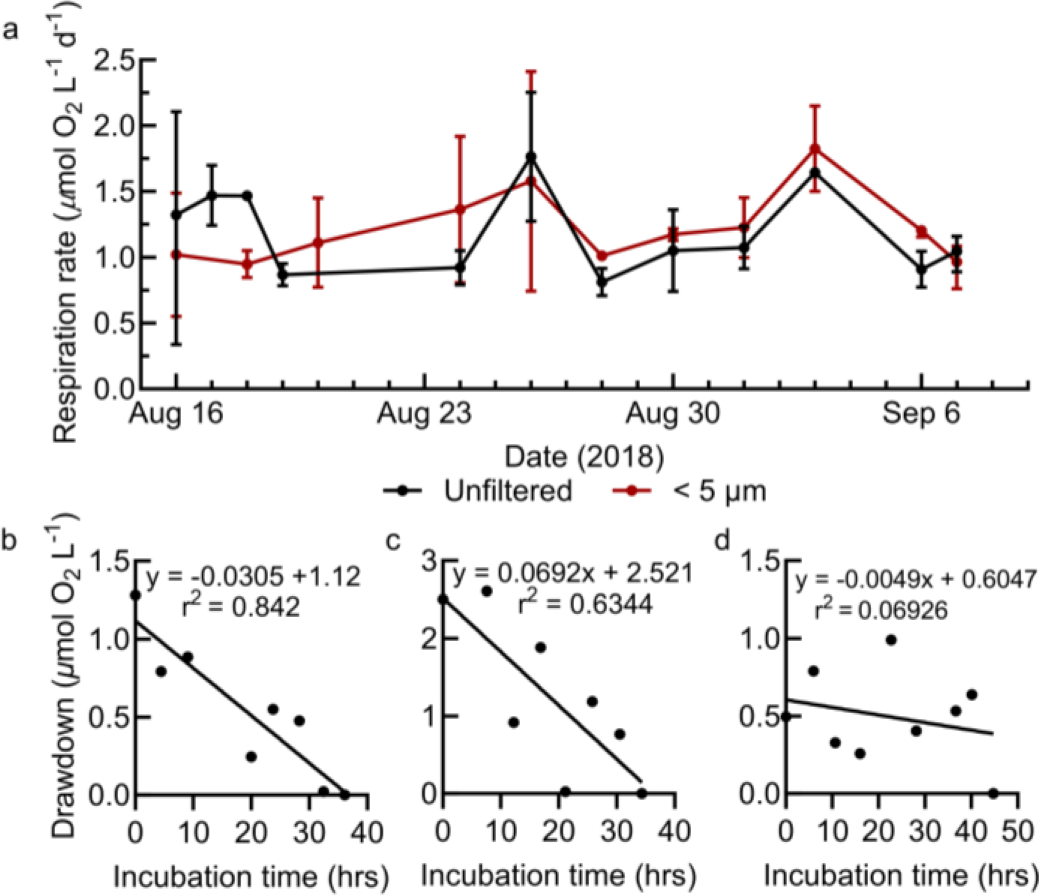
(a) Range of surface respiration rates for the unfiltered and 5 *µ*m size fractions during the N. Pacific sampling. (b-d) Select N. Pacific respiration assays representative of strong, moderate, and weak fits of O2 drawdown. Model I linear regression statistics are shown in upper right. All N. Pacific O2 drawdown assays are shown in Supplemental Figure 1.

The unfiltered and small size fraction respiration rates were often statistically indistinguishable in the N. Pacific (Fig. 2 and 3). This may be due to the combination of optode precision, the low frequency manually obtained measurements, and low O2 drawdown. As discussed by del Giorgio and Williams (2005), the magnitude of the O2 signal is a particularly influential driver of uncertainty in low activity environments such as this region. This methodological limitation explains instances where the derived small fraction respiration exceeds the unfiltered rates, though still within the error of biological replication. In general, though, these results suggest that the majority of microbial community respiration in the mixed layer is attributable to small cells <5 *µ*m, likely consisting of bacterioplankton, cyanobacteria, and small eukaryotes (McNair et al. 2021; Meyer et al. 2022).

**Figure 3.**
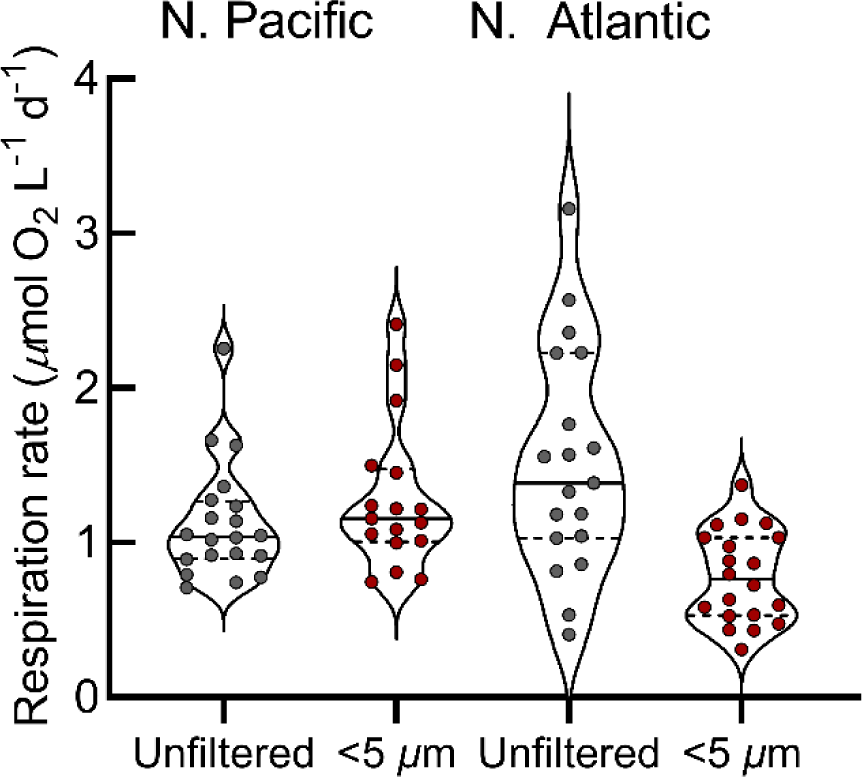
Mixed layer respiration rates across the N. Pacific and N. Atlantic provinces. Violin plots show respiration rates with the median (solid horizontal lines) and the 25^th^ and 75^th^ percentiles (dashed horizontal lines); unfiltered (gray points), filtered (red points).

Vertical profiles from the N. Pacific show unfiltered respiration rates decreasing from the surface to 50 m, before increasing to a near maximum at 95 m, and then decreasing to below reliable quantification at 200 m (Fig. 4, S1). Bacterial production did not increase at 95 m, in contrast to the relatively high respiration here. Instead, BP steadily decreases from the surface to depth (Fig. 4; Stephens et al. 2023).

**Figure 4.**
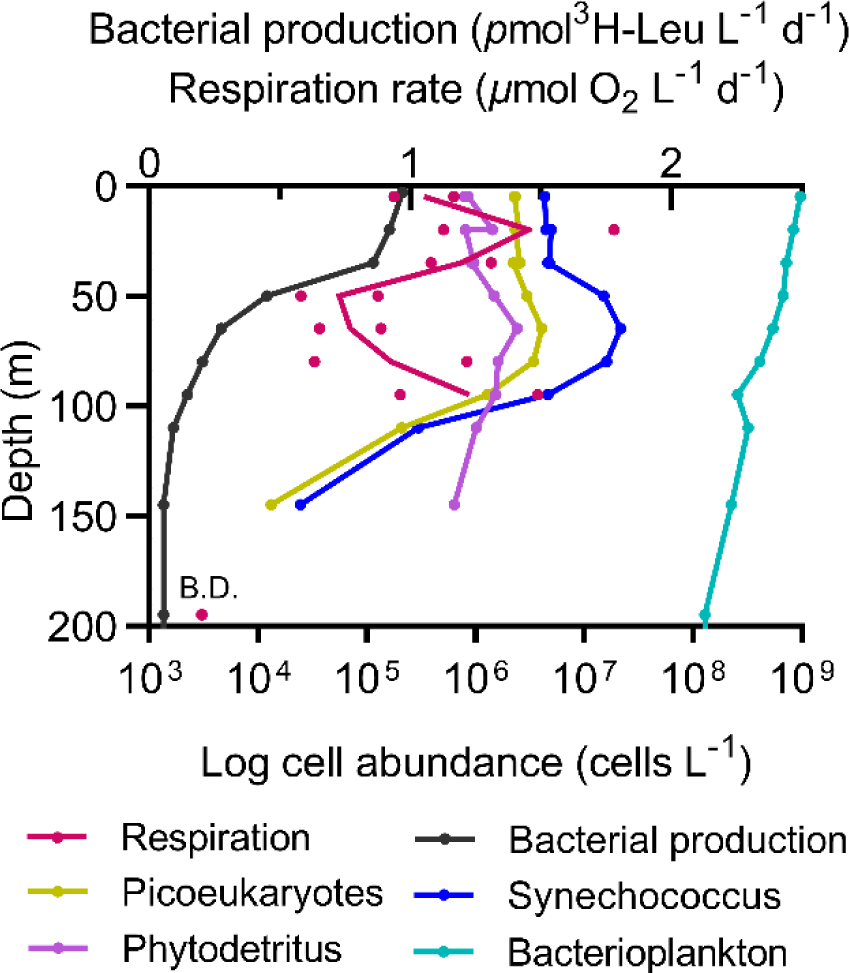
N. Pacific vertical unfiltered respiration rates, bacterial production, phytoplankton abundances, and phytodetritus on September 3, 2018 at Ocean Station Papa.

### N. Atlantic respiration rate assays

Although we were able to estimate respiration rates and determine that small cells contribute the most to community respiration in the N. Pacific, we were limited in our ability to robustly resolve trends and variations among size fractions due to the low frequency of manually obtained O2 measurements. To improve on this, in the N. Atlantic at the PAP site (Fig. 1), we increased the number of oxygen meter channels and directly mounted the polymer optical fibers to the BOD bottles (Fig. S2), enabling simultaneous O2 measurements every 30 seconds for 12 sample bottles and one Milli-Q control. As a result, the number of O2 measurements increased from *c.a.* ten in the Pacific to *c.a.* 3,000 measurements per assay in the Atlantic (Fig. 5a). The high-frequency O2 measurement system employed reduced the need for time-consuming, manually acquired measurements, increased the accuracy of respiration rate determinations, and reduced the incubation time needed to obtain a detectable signal. The high resolution O2 measurements enabled clearly identifying which parts of the oxygen drawdown curve needed to be trimmed from the start, as well as determining whether the curve became non-linear towards the end of the experiment and required end trimming. A rolling mean was applied to the large number of point measurements, which was then used to calculate the rate (Fig. 5a). These improvements allowed us to calculate O2 consumption rates with greater accuracy than the point resolution of the instrumentation (± 1.4 *µ*mol L^-1^ O2 at 283 *µ*mol L^-1^ O2; OXY4-SMA, PreSens), and reduce the incubation time required to obtain a reliable drawdown signal in this environment to under 20 h. With this approach, we were able to resolve respiration rates <0.2 *µ*mol O2 L^-1^ d^-1^ (Fig. 5d-i, S3).

**Figure 5.**
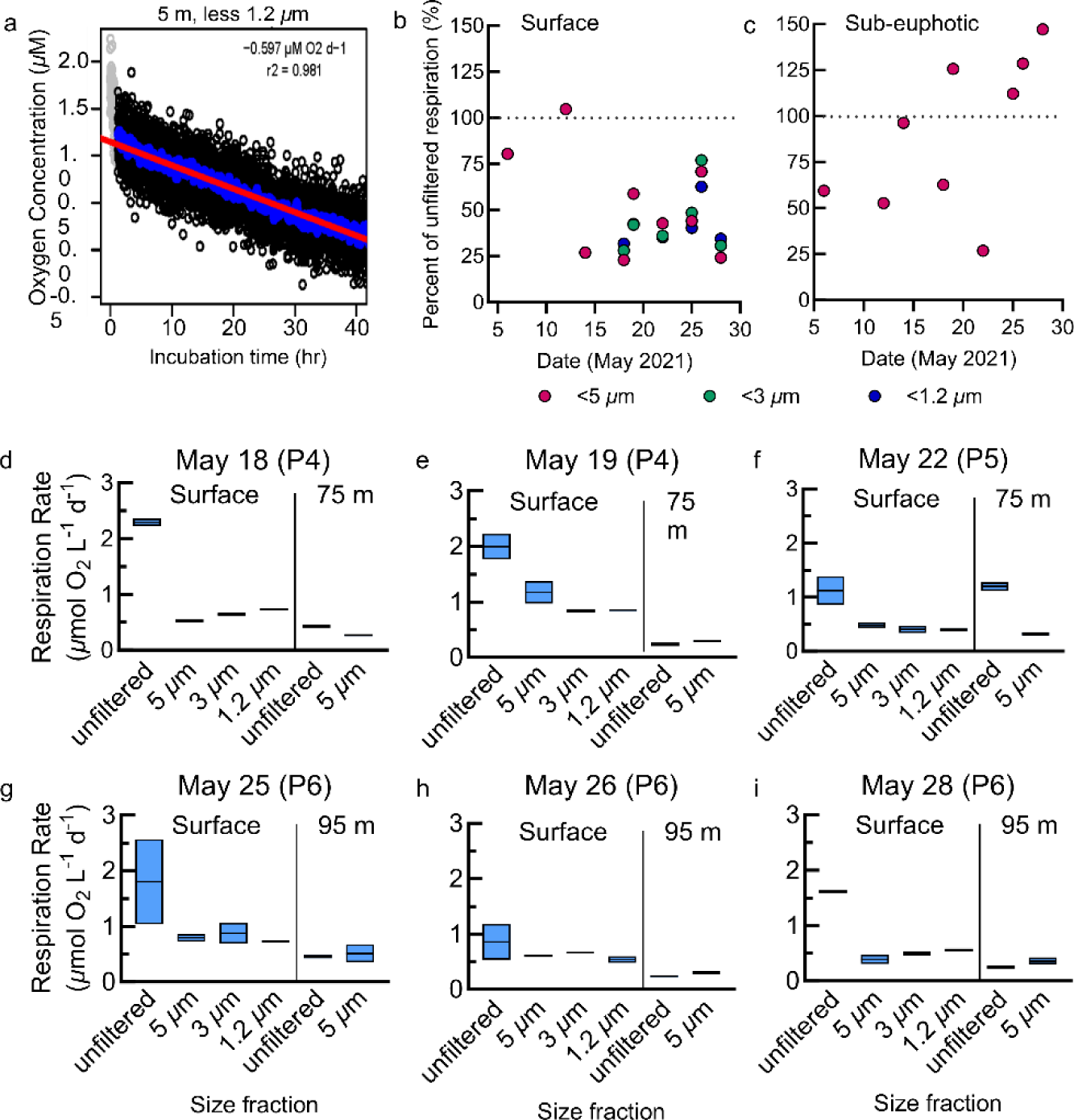
(a) Representative O2 drawdown assay from the N. Atlantic showing individual O2 measurements made every 30 s (black circles). Gray points represent sample points determined to be in the acclimation period and are excluded from calculations. The linear regression (red line) is calculated from the 12.5-min rolling average (blue circles). Plots for all N. Atlantic assays are in Supplemental Figure S3. (b-c) The percent contribution of each size class to total (unfiltered) respiration at the surface (b) and sub-euphotic zones (c) in the N. Atlantic. The size classes are nested rates, where the <3 *µ*m fraction includes respiration from the <1.2 *µ*m fraction, and so forth. (d-i) Size-fractionated respiration rates in the N. Atlantic. Assays of unfiltered, small <5 *µ*m, micro <3 *µm*, and nano <1.2 *µ*m filtrate were run in duplicate at the surface and below the euphotic zone beginning May 18^th^ with the storm period noted following the date of each plot. The boxes represent the range and average of the duplicates.

### Comparison to INT rates

To validate the accuracy of the optode assays, we compared them to several INT derived measurements during the N. Atlantic campaign. The INT respiration method relies on a colorimetric change resulting from the oxidation of the coenzyme Q-cytochrome B complex step in the respiratory electron transport system of bacteria, phytoplankton, and zooplankton (Martínez-García et al. 2009; García-Martín et al. 2019). The most suitable comparison between size fractions collected across these approaches involves the optode <3 *µ*m fraction and the INT 3 to 0.22 *µ*m fraction. The INT respiration rates ranged from 0.55 to 0.80 *µ*mol O2 L^-1^ d^-1^, closely aligning with our optode ranges from the same dates (0.42 to 0.84 *µ*mol O2 L^-1^ d^-1^) (Fig. 6). The consistency between the INT and optode-based methods supports the accuracy of both assays. García-Martín et al. (2019) conducted an extensive validation of INT-based respiration rates compared to oxygen drawdown-based rates, also finding reasonable agreement between the methods.

**Figure 6.**
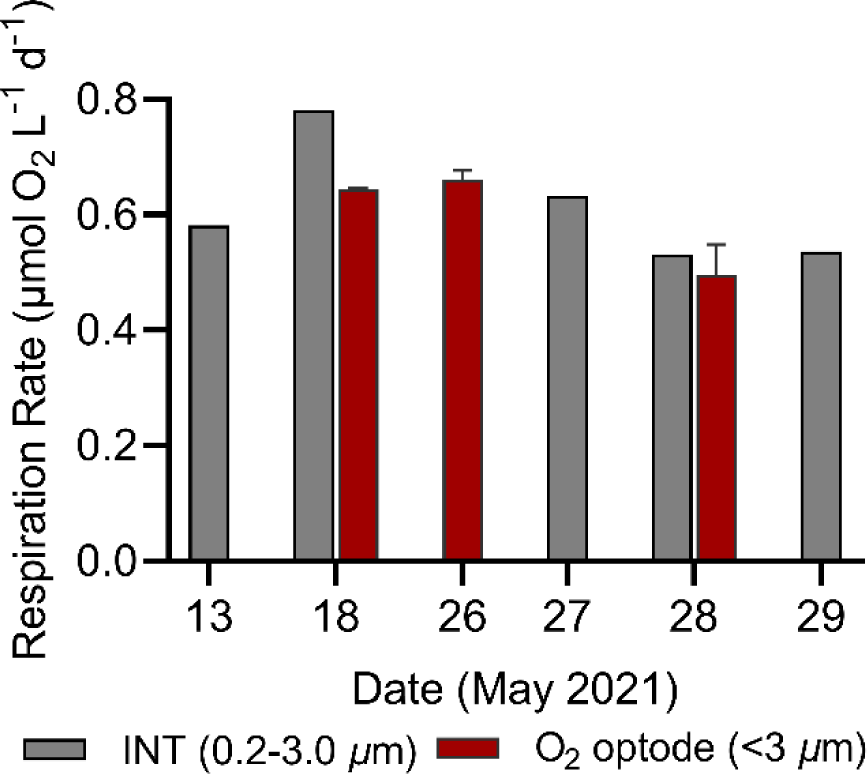
Comparison of surface *in vivo* 2-para-(iodophenyl)-3(nitrophenyl)-5(phenyl) tetrazolium chloride (INT, 0.2-3.0 *µ*m) and O2 optode derived (<3 *µ*m) respiration rates for several dates in the N. Atlantic. Error bars represent standard deviations of optode duplicates.

### Potential DOC enrichment

Organic carbon contamination can be a significant challenge in obtaining accurate respiration rates in oligotrophic systems, considering the low *in situ* labile DOC concentrations and the general carbon limitation of heterotrophic marine bacteria. During sample handling and incubation setup, measures were taken to minimize carbon contamination by adhering to the practices outlined above. Additionally, TOC and DOC concentrations were assessed in several samples during the N. Atlantic campaign. The average *in situ* TOC concentration was 58.5 µmol C L^-1^ at 5 m and 54.9 µmol C L^-1^ at 75 m with daily variability of 1-3 µmol C L^-1^. A modest increase in TOC concentration (up to 6 to 8 *µ*mol C L^-1^) and DOC (up to 9 to 11 *µ*mol C L^-1^) was observed in the BOD bottle incubations compared to samples collected directly from the cast (Fig. S4). However, most TOC comparisons revealed a difference of 2 to 3 *µ*mol C L^-1^, and consistent replication of respiration rates across the BODs suggests that the enrichment was not all bioavailable over the time of incubation.

### Size class contributions

Size-fractionated respiration assays (unfiltered and small fraction (<5 *µ*m) were conducted at the surface and just below the euphotic zone in the N. Atlantic. For six surface samples, we included two additional size classes: micro, and nano (Fig. 5d-i). The *unfiltered* fraction represents the total microbial community; the *small* (<5 *µ*m) contains large particles, phytoplankton, and bacterioplankton; *micro* (<3 *µ*m) contains pico-eukaryotes, particle-associated, and free-living bacteria; and the *nano* (<1.2 *µ*m) fraction contains free-living bacteria and some pico-eukaryotes.

PAP surface respiration rates averaged 1.7 *µ*mol O2 L^-1^ d^-1^ for the unfiltered and 0.9 *µ*mol O2 L^-1^ d^-1^ for the small size fraction (Fig. 5b). The small, micro, and nano classes averaged 45% of total respiration at the surface (Fig. 5b, c), with differences between these classes often not statistically distinguishable (Fig. d-i). Since the filtered fractions represent nested respiration rates, equal rates of the smallest two size fractions suggests that the <1.2 *µ*m community is contributing the most to respiration rates. This is consistent with Aranguren-Gassis et al. (2012) who showed surface bacterial respiration contributed approximately 30% of community respiration across a range of ocean systems. Similarly, Fasham et al. (1999) modeled community member contributions to respiration in the Northeast Atlantic near PAP in May 1990 and report 38% contribution by bacteria, 30% by phytoplankton, and 57% by the <5 *µ*m fraction.

N. Atlantic respiration rates were up to 10 times lower below the euphotic zone (either 75 m or 95 m), averaging 0.6 *µ*mol O2 L^-1^ d^-1^ for unfiltered fractions and 0.3 *µ*mol O2 L^-1^ d^-1^ for small fractions (Fig. 5 and 7). A notable disparity between the surface and sub-euphotic was clear in size class contributions to total microbial respiration. The small fraction constituted an average 53% of total respiration at the surface, as opposed to 90% in the sub-euphotic (Fig. 5c), implying a shift in the predominant active microbial community classes below the euphotic zone. Note the micro- and nano-respiration fractions were not collected below the euphotic zone due to limitations in available sensors.

**Figure 7.**
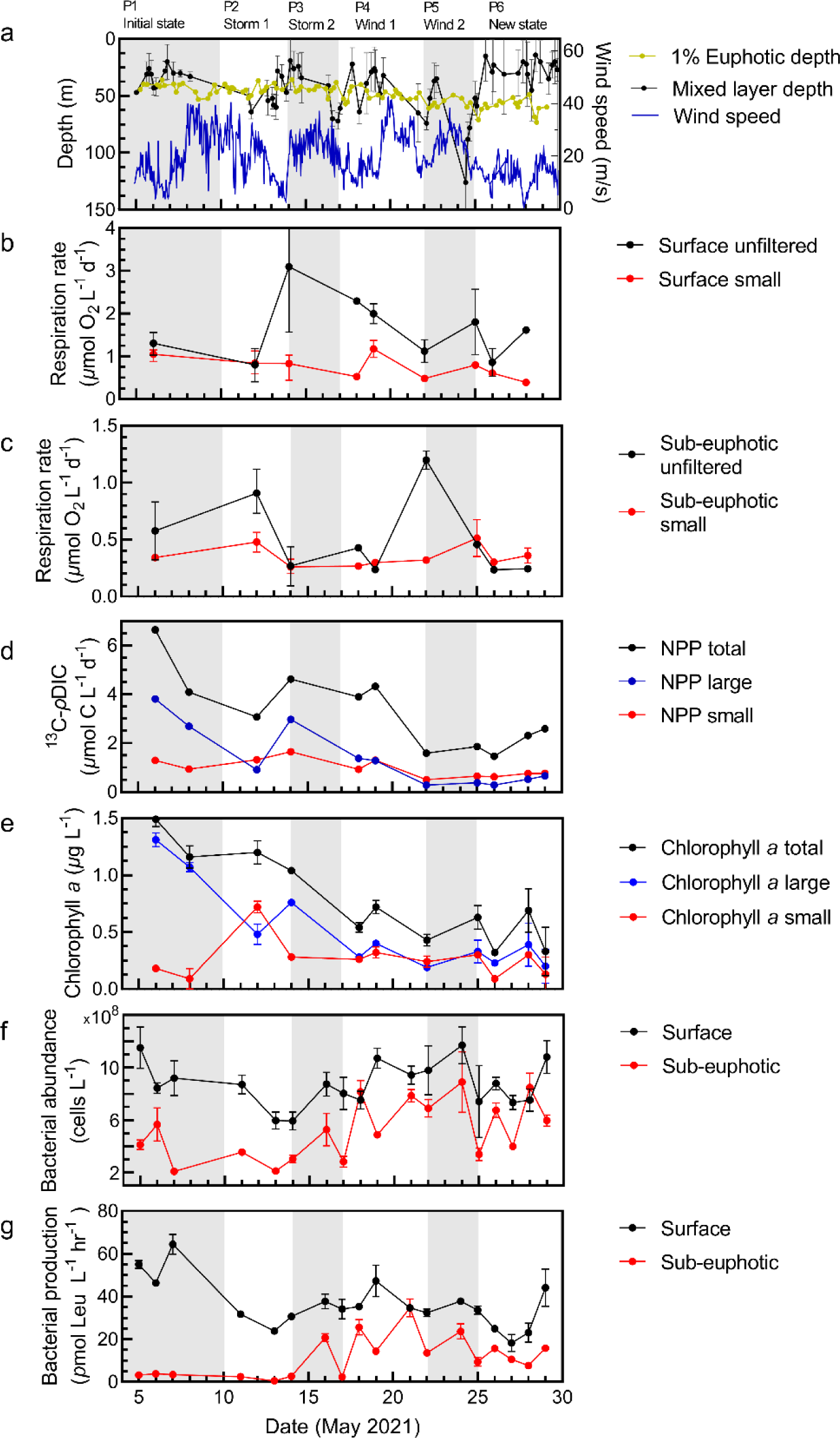
N. Atlantic physical context, respiration, productivity, and secondary production rates at the Porcupine Abyssal Plain at the surface (5 m) and sub-euphotic depths (75 m or 95 m). Gray and white shading indicates the periods for arrival on station (P1), storm (wind and cloud cover/precipitation) or wind-only events and recovery (P2-P5), and new system state (P6). Physical parameters include mixed layer depth and euphotic zone depth (1% Io) (a). Size-fractionated microbial respiration mean and range at the surface (b) and below the euphotic zone (c) for the unfiltered and <5 *µ*m fractions. Mean and standard deviations for surface chlorophyll *a* concentration (d) and net primary production (e) are shown for the total, small, and large fractions. Bacterial abundance (f) and production mean and standard deviations (g) at the surface and sub-euphotic zone match the sampling depths for respiration.

### Replicate variability among size fractions and depths

Variation between replicate respiration assays demonstrated a dependency on both depth and size fraction. A residual standard error (RSE) analysis indicated that the mean error was low for surface, size-fractionated classes but significantly higher for the unfiltered fraction (Dunnett’s T3 multiple comparisons test, *p* <0.001). Additionally, surface unfiltered replicates exhibited higher coefficients of variation (66.2% CV, *F*5.000, 27.38* = 16.65) compared to the fractionated assays (23% small, 26% micro, 16% nano.; Fig. S5, S6). In contrast, no significant differences in residual errors and % CV were observed for sub-euphotic samples, with the percentage of outliers decreasing similarly with size fractions (Fig. S5, S6). The increased variability in large and unfiltered samples likely stems from the stochasticity in capturing large plankton and particles present at lower concentrations in the samples.

### Temporal respiration responses to mixing events

In contrast to the relatively stable N. Pacific Station Papa system where primary production, export, and physical mixing remained fairly constant, the N. Atlantic PAP survey exhibited greater dynamics, encompassing a phytoplankton bloom decline and multiple storms and associated mixing events (Fig. 7). Consequently, we divided the temporal trends of the N. Atlantic campaign into six periods (P1-6): the initial system state, four mixing events, and the final system state.

The initial system state at PAP in May was characterized by a declining large (>5 *µ*m) predominately diatom phytoplankton bloom, exhibiting a decrease in both NPP and chlorophyll *a* (1.5 to 0.3 *µ*g Chl *a* L^-1^) (Fig. 7d-e). Surface microbial respiration rates were elevated in the small fraction, indicative of the high initial phytoplankton and bacterioplankton cell abundances. A storm event (start of P2) then deepened the mixed layer, decreasing surface stocks and rates of primary producers and microbial respiration. However, below the photic zone, deep unfiltered respiration rates increased sharply. This switch reflects a redistribution of biomass in the water column by the storm, with large phytoplankton from the surface driven deeper (Fig. 7b-d; Meyer et al. 2023). With time, the surface respiration rates increased again to reach an observed maximum.

A second storm event (P3) deepened the mixed layer again and there was another reduction in primary production, stocks, and surface respiration into P4 (Fig. 7a-e). During P3, the phytoplankton shifted to a predominantly haptophyte and dinoflagellate community (Meyer et al. 2023; https://ifcb-data.whoi.edu/timeline?dataset=EXPORTS) and the ratio of large to small-cell phytoplankton began to equalize (Fig. 7e). Deeper, bacterial abundance, and notably BP, increased (Fig. 7f, g).

A period of intense and sustained wind events (P4 and P5) caused the mixed layer to reach new deep depth of 120 m, before shoaling rapidly to 20 m (Fig. 7a). We were able to incorporate additional respiration size classes into surface sampling efforts for P4 and P5. The contribution of the small fraction relative to the micro- and nano-fractions decreased in P4 and P5 until the three fractions were indistinguishable by P6 (Fig. 5), indicating a shift in microbial community structure (Fig. 5). This increased contribution of small cells to total respiration corresponds with increased bacterial abundance (Fig. 5b; 7b, f). Notably, a cruise maximum in unfiltered sample respiration rates below the euphotic zone of 1.2 *µ*mol O2 L^-1^ d^-1^ is observed at the conclusion of P4, aligning with the timing of a major flux event (Fig.7c) and increased particulate organic carbon abundance over the duration of the sampling (S Clevenger pers. comms.).

During the final observation period (P6), the mixed layer was relatively stable and shallow, and the community was in a new state characterized by increasing net primary production and BP (Fig. 7d, g). Surface unfiltered respiration exhibited elevated levels, whereas the small fraction maintained a stable trend, possibly due to higher particle-associated respiration towards the end of the cruise and low activity of the phytoplankton (Fig. 7b, d, e). Below the euphotic zone, particulate organic carbon was high (Estapa et al. 2023; S Clevenger pers. comms.) and unfiltered respiration rates and filtered rates were similar in magnitude (*c.a.* 0.3 *µ*mol O2 L^-1^ d^-1^, Fig. 7c).

In summary, the temporal and vertical trends during the N. Atlantic sampling revealed that surface respiration rates peaked midway through the campaign, and then decreased, correlating with BP (Pearson’s correlation, *r*=0.83, *p*=0.01; Fig. S7). However, periodic storm events disrupted this pattern by redistributing biomass from the surface to deeper waters, diluting the surface community while simultaneously promoting increased respiration and BP within the deeper community. Respiration trends at the beginning of the cruise (P1-P4) correspond to the redistribution of phytoplankton biomass. As the bloom biomass declined and shifted from diatom to dinoflagellate and haptophyte dominated community state, large cell respiration was more strongly influenced by the formation and flux of fluffy particles which distributed vertically based on wind events and shifting mixed layer depth (C Durkin pers. comm.; S Clevenger pers. comms.). The flux of particles appears to have stimulated small cell respiration at depth, supported by a moderate but significant relationship between large-cell chlorophyll *a* with sub-euphotic BP (Pearson’s correlation, *r*=0.38, *p*=0.04; Fig. S7) supporting the idea of heterotrophic utilization of sinking particles while increased bacterial abundance led to greater bacterial contribution to total respiration at both depths.

### Diel variability

It is increasingly recognized that marine microbial communities have diel metabolic rhythms driven in part by phytoplankton release of highly labile dissolved organic matter (Gifford et al. 2014; Ottesen et al. 2014; Becker et al. 2018). Given we observed a significant relationship between small cell NPP and small cell respiration (Pearson’s correlation, *r*=0.68, *p*=0.05; Fig. S7), we hypothesized diel primary production patterns may drive diel fluctuations in respiration. To test this, we collected unfiltered seawater and incubated it under ambient light and temperature conditions to replicate phytoplankton diel activities and dissolved organic matter production. On-deck incubations were periodically sampled every 6 h for respiration assays of the nano (<1.2 µm) community to observe potential diel responses to dissolved organic matter changes. We observed no statistical differences in respiration rates over the 42-h period (Fig. S8).

There are two potential explanations for the absence of a diel respiration trend. One possibility is that respiration rates genuinely remain constant throughout the diel cycle, as observed in other studies. Robinson et al. (2019) found that bacterial carbon demand in the Celtic Sea did not change over diel cycles using Winkler titration and INT-based approaches. Similarly, Martinez-Garcia and Karl (2015) examined microbial community respiration every 4 h over a 44-h period in the Subtropical N. Pacific using the INT approach and discovered no trends between time points or significant differences between night and day rates. Alternatively, our observed lack of a diel response could be attributed to the rapid utilization of any highly labile signal within the first 2.5 h of incubation, which would be undetectable due to our optode sensor spot acclimation period. If this is the case, it represents an important methodological limitation, as the rapid consumption of highly labile carbon by the bacterial community would not be captured, resulting in an underestimation of respiration and derived bacterial carbon demand.

## SUMMARY

In this study, we evaluated microbial respiration rates across various size classes and depth horizons by utilizing O2 drawdown assays with optode measurements. Measuring accurate respiration rates has traditionally been a difficult task, especially in oligotrophic regions due to low rates which challenge the precision of current optical techniques (Robinson 2019). However, here, by attaching the optical fiber directly to the BOD bottle and performing frequent measurements in a stable temperature environment, the assay’s quantitative capabilities were considerably improved. These enhancements increased the accuracy and robustness of measuring O2 drawdown in the surface and upper mesopelagic zones of the oligotrophic ocean. The measured respiration rates aligned well with both INT and historical measurements. The resolution of the continuous-mounted optical sensor method and the capacity to resolve rates as low as 0.2 *µ*mol O2 L^-1^ d^-1^ provides a means to expand respiration rate coverage and precision in the open ocean. This contributes to filling gaps in datasets and creating a more tightly constrained carbon budget, which is crucial for understanding global carbon cycling.

However, improvements are needed to enhance the accuracy and utility of the technique further. Many microbial processes occur on hourly or shorter time scales (Ottesen et al. 2014; Becker et al. 2018; Muratore et al. 2022), which are currently not detectable with optode incubation assays due to the sensor’s adaptation period. This limitation might be one reason we did not detect diel respiration trends during our N. Atlantic diel experiment. Additionally, BP assays, which provide the complementary measurement required to calculate bacterial carbon demand and close the microbial carbon budget, are typically measured on much shorter timescales (several hours in total) (Baetge et al. 2021). This could result in a temporal disconnect between these two critical carbon cycle measurements. Yet, the close correlation observed in our study between the rates obtained from INT, which has a relatively brief incubation time of 2 h, and those from the optode assays, with a longer duration ranging from 20 to 48 h, suggests that the extended incubation periods and initial acclimation phase of the optode assays did not significantly impact the rates compared to shorter-term incubation methodologies in this study. Stephens et al. (2023) reported a similar agreement between optode and short-term BP assays. Future improvements to optode-derived O2 incubation assays should aim to minimize the spot adaptation period and ideally enhance spot sensitivity.

Utilizing these optode assays in the N. Pacific and N. Atlantic, we observed varying contributions of different microbial size classes to community respiration in these two contrasting oceanic provinces. The N. Pacific sampling at Station Papa represented the low productivity endmember where our measured rates were relatively low (average 0.84 *µ*mol CO2 L^-1^ d^-1^ using a respiratory quotient of 1.4 O2:CO2) but appeared elevated compared to derived respiration rates from BP assays (0.19 *µ*mol CO2 L^-1^ d^-1^; Stephens et al. 2020). It is possible that the manually obtained, low temporal resolution measurements of the N. Pacific assays could not accurately constrain these low rates. However, it is important to note that our small size fraction encompasses a significantly larger portion of the microbial community than that of the <1 *µ*m fraction used for BP assays and that our rates are similar to historical measurements by Sherry et al. (1999). The <5 *µ*m fraction included not only heterotrophic bacteria but also abundant phototrophic *Synechococcus*, picoeukaryotes, and small grazers, which may all contribute substantially to community respiration (Langdon, 1993; Marra and Barber 2004; D Steinberg, and A Maas pers. comm.).

Both biomass and rate measurements in the N. Pacific were dominated by small cells (Meyer et al. 2022). Small cell (<5 *µ*m) respiration rates were often statistically indistinguishable from the whole water samples (Fig. 2a, 3). Linear regressions show that respiration positively correlated with small cell phytoplankton biomass (*r^2^*=0.50), weakly correlated to coinciding program measurements of particulate organic carbon (*r^2^*=0.21) and exhibited an inverse relationship with NPP. These findings reflect the low biomass, low productivity, and highly recycled nature of the Station Papa ecosystem in late summer (Meyer et al. 2022; McNair et al. 2023).

Below the mixed layer, the N. Pacific vertical profiles of respiration exhibit an expected decrease with depth, consistent with minimal particle flux occurring at this time (Estapa et al. 2021). However, respiration rebounds just below the deep chlorophyll maximum. Interestingly, there is no concurrent increase in BP or bacterial abundance observed at the base of the deep chlorophyll maximum. In fact, below 65 m, respiration displays an inverse relationship with BP and small phytoplankton abundances. This pattern suggests a decoupling of microbial respiration from production, biomass, and abundances. A midwater increase in respiration rates and decoupling with BP have been observed elsewhere as discussed by Martina-Garcia and Karl (2015) who also saw respiration rates in the subtropical N. Pacific with similar mid-water respiration maximum. Additionally, Kim et al. (2017) found a decoupling between BP and bacterial respiration in downwelling systems where nutrients are low, proposing that a shift in metabolism towards cellular maintenance rather than growth was driving decoupling.

During our sampling at Station Papa, we postulate that bacteria around the 100 m depth horizon were utilizing phytodetritus, leading to an increase in respiration despite the divergence from the BP trend. The 95 m respiration increase is just below the base of the euphotic zone (1% PAR, Siegel et al. 2021), the deep chlorophyll maximum, and the phytodetritus maximum which presumably originates from high densities of cyanobacteria and picoeukaryotes (Fig. 4). Simultaneous work by Stephens et al. (2020) shows a decrease in bacterial growth efficiency with depth here, indicating a higher proportion of respiration to production. Furthermore, the abundant, *Synechococcus* and picoeukaryotic communities may not be well captured by ^3^H leucine-based BP estimates due to the inherently flexible metabolism of these mixotrophs and the dependance of their secondary production on light availability photosystem activity (Duhamel et al. 2018). Below 100 m, respiration, BP, bacterial abundance, and phytodetritus concentrations decreased to their lowest values.

N. Atlantic sampling at PAP showed a higher activity environment with substantial system dynamics. Here, larger phytoplankton were much more prevalent, resulting in periods in which large cell primary production exceeded that of the small (Meyer et al. 2023). Accordingly, our N. Atlantic respiration rates in the unfiltered fraction were on average 1.4-fold higher than that observed in the N. Pacific. The small cell respiration rates, however, were on average 1.3-fold lower in the N. Atlantic than in the N. Pacific and matched with BP derived bacterial respiration estimates as well as several overlapping INT-determined rates. The lower small size fraction rates may be due to the technical improvements used in the N. Atlantic (increasing temporal measurement resolution to >1000 points), and or a relatively smaller contribution of <5 to <1 *µ*m cell sizes in the N. Atlantic versus N. Pacific.

On average, the small cell respiration in the N. Atlantic was responsible for 45% of the total respiration, within 17% of prior estimates for this system (Fasham et al. 1999; Aranguren-Gassis et al. 2012; Binetti et al. 2020). The contribution of the size classes to community respiration was greatest in the mid- to late stages of the observation period as phytoplankton biomass in the surface decreased. Further size fractionation allowed us to more finely resolve microbial size class contributions to community respiration. After May 22^nd^, these three smallest size fractions displayed little difference in magnitude at the surface, coinciding with the second bloom phase where phytoplankton biomass had shifted from large to small cells (Meyer et al. 2023). This suggests that the smallest fraction, <1.2 *µ*m, was contributing the most to filtered respiration rates.

The N. Atlantic system exhibited substantial respiration dynamics driven by the decline of a phytoplankton bloom and multiple significant mixing events. As the bloom dissipated, respiration trends were initially high when phytoplankton biomass was high, affecting the entire upper water column due to the redistribution of biomass caused by storm events. Respiration typically returned to pre-storm levels after storms. When the bloom was no longer dominated by large diatoms, respiration was high where particles were present, and small cell respiration accounted for a larger portion of community respiration. Below the euphotic zone, respiration rates were much lower, with a significantly higher contribution (80%) of the small size fraction to total respiration. Similar dynamics in the N. Atlantic during spring blooms have been documented by the Biogeochemical Ocean Flux Study Spring Bloom Experiment in 1990, the N. Atlantic Bloom Experiment, and recent research near PAP (Bender et al. 1992; Savidge et al. 1992; Binetti et al. 2020), all of which demonstrate that this is an energetic system, net autotrophic annually, with frequent switching between net autotrophic and heterotrophic states. Comparatively, Bateage et al. (2022) also captured a restratification event in the western N. Atlantic where microbial activity increased as POC and chlorophyl *a* decreased, suggesting solubilization and partial consumption of redistributed particulate matter. Our findings reveal that these dynamics can occur on relatively short timescales (days) and drive interactions between respiration rates in the surface and euphotic zones.

Our findings provide insights into microbial carbon cycling and export potential in these two distinct ocean provinces. In the N. Pacific at Station Papa, we estimate a mean 58% (± 20%) of GPP is respired at the surface. When accounting for all other measured food web respiration components, approximately 100% of GPP is estimated to be respired, resulting in the low carbon export potential of this system observed during our sampling (Buesseler et al. 2020; McNair et al. 2023). In the N. Atlantic at PAP, we estimate that 34% (± 25%) of GPP is respired at the surface. Though this represents a preliminary, minimum bound for N. Atlantic food web respiration, production surpasses respiration here, supported by high coinciding particle flux, leading to relatively more carbon export from the mixed layer compared to the N. Pacific (Estapa et al. 2023; S Clevenger pers. comms.).

While our data were gathered from two well-studied oceanographic stations over several weeks, more extensive sampling is required across both temporal and spatial scales to better resolve carbon cycle components. Furthermore, our findings reveal that significant fluctuations in respiration rates can occur at daily time scales, which can considerably affect estimates of net community production within the system. Facilitated in part by advancements in respiration measurement techniques, like those detailed here, increased respiration measurements will enable a more accurate assessment of respiration’s role in ocean ecosystem metabolism.

## Author Contributions

This work was conceptualized by SG and MRC with support from CAC, BS, AM, and NC. MRC and SG were lead investigators and administrators for this work with support from all authors. Supervision was provided by SG (project lead), JRG (a program lead), and CAC (supporting). Data curation, methodology, formal analysis, and visualizations were completed by MRC and SG (leads) with INT, BP, and TOC led by BS, and CAC; productivity by MGM, and POC and phytoplankton abundances by JRG. Pacific respiration assays were conducted by SG (lead) supported by GCS. National Aeronautics and Space Administration (NASA) Exports funding was awarded to AM, SG, NC (equal) and CAC (supporting) with resources primarily supplied by SG. The work was drafted by MRC and SG (leads) with support from BS and MGM. All authors reviewed and approved the manuscript.

## Acknowledgements

The authors thank the NASA EXPORTS team including program, leads D. Seigel and I. Cetinić, chief scientists D. Steinberg and J. Graff, and the 2018 and 2021 EXPORTS team. We thank the captain and crew of the R/V *Roger Revelle* (RR1813) and RSS *James Cook* (JC214) for their facilities, expertise, and dedication. AM, SG, and NC received National Aeronautics and Space Administration award 80NSSC17K0552 and award 80NSSC18K0437 to CAC. We declare no conflicts of interest.

## Data Availability Statement

All EXPORTS data, metadata, and methods referenced herein are available on the National Aeronautics and Space Administration (NASA’s) data archive at seabass.gsfc.nasa.gov; DOI: 10.5067/SeaBASS/EXPORTS/DATA001. N. Atlantic cruise JC214 particulate organic carbon data can be found on SeaBASS under UCSB/CRSEO/EXPORTS/EXPORTSNA/archive. DOC, 3H-Leucine, and bacterial abundance can be found on SeaBASS under UCSB/carlson/EXPORTS/EXPORTSNA/archive.

