## Supplemental Information for "Microbial Respiration in Contrasting Ocean Provinces via High-Frequency Optical Assays"

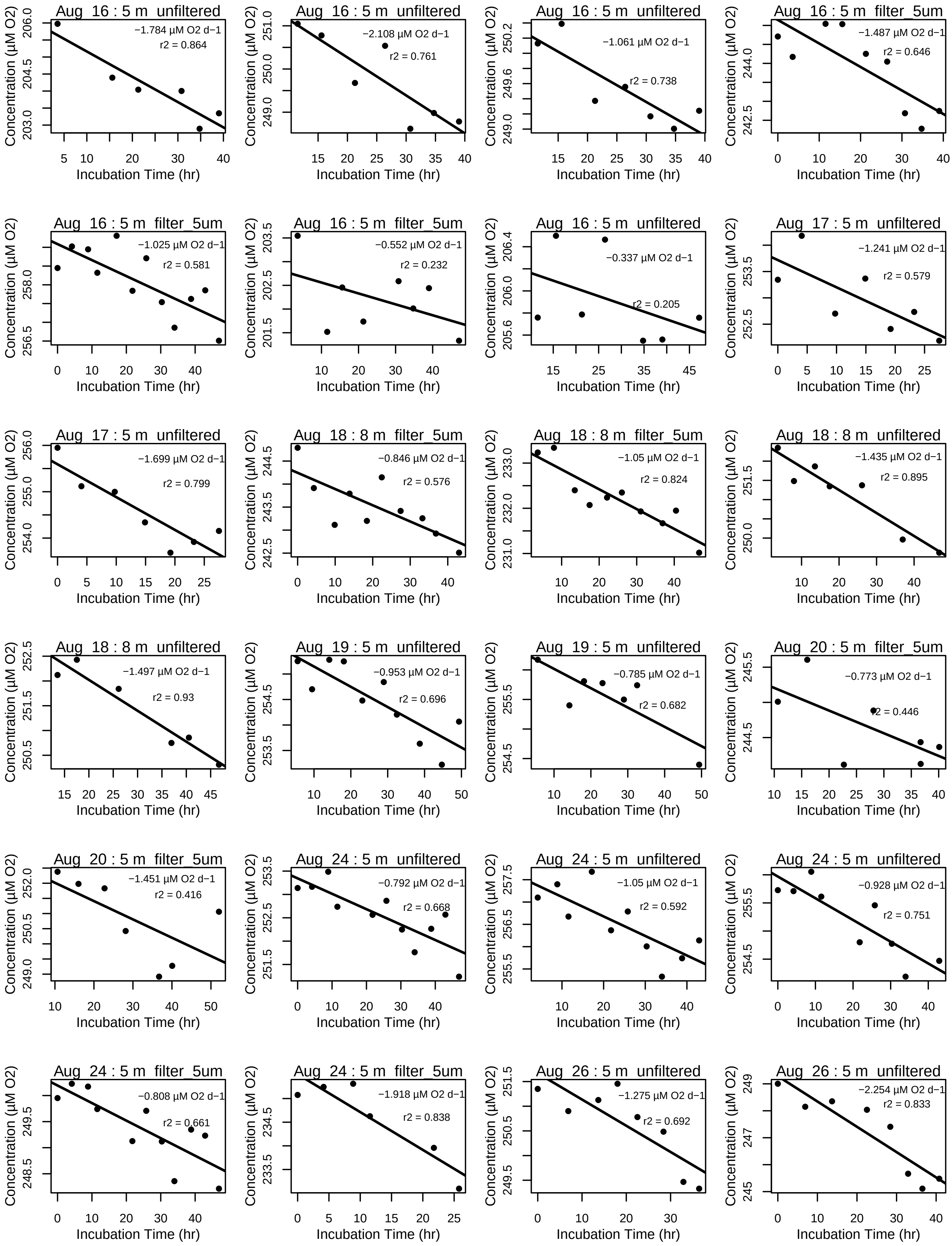
Aug 16 : 5 m unfiltered
−1.784 µM O2 d−1
r2 = 0.864
Aug 16 : 5 m unfiltered
−2.108 µM O2 d−1
r2 = 0.761
Aug 16 : 5 m unfiltered
−1.061 µM O2 d−1
r2 = 0.738
Aug 16 : 5 m filter_5um
−1.487 µM O2 d−1
r2 = 0.646
Aug 16 : 5 m filter_5um
−1.025 µM O2 d−1
r2 = 0.581
Aug 16 : 5 m filter_5um
−0.552 µM O2 d−1
r2 = 0.232
Aug 16 : 5 m unfiltered
−0.337 µM O2 d−1
r2 = 0.205
Aug 17 : 5 m unfiltered
−1.241 µM O2 d−1
r2 = 0.579
Aug 17 : 5 m unfiltered
−1.699 µM O2 d−1
r2 = 0.799
Aug 18 : 8 m filter_5um
−0.846 µM O2 d−1
r2 = 0.576
Aug 18 : 8 m filter_5um
−1.05 µM O2 d−1
r2 = 0.824
Aug 18 : 8 m unfiltered
−1.435 µM O2 d−1
r2 = 0.895
Aug 18 : 8 m unfiltered
−1.497 µM O2 d−1
r2 = 0.93
Aug 19 : 5 m unfiltered
−0.953 µM O2 d−1
r2 = 0.696
Aug 19 : 5 m unfiltered
−0.785 µM O2 d−1
r2 = 0.682
Aug 20 : 5 m filter_5um
−0.773 µM O2 d−1
r2 = 0.446
Aug 20 : 5 m filter_5um
−1.451 µM O2 d−1
r2 = 0.416
Aug 24 : 5 m unfiltered
−0.792 µM O2 d−1
r2 = 0.668
Aug 24 : 5 m unfiltered
−1.05 µM O2 d−1
r2 = 0.592
Aug 24 : 5 m unfiltered
−0.928 µM O2 d−1
r2 = 0.751
Aug 24 : 5 m filter_5um
−0.808 µM O2 d−1
r2 = 0.661
Aug 24 : 5 m filter_5um
−1.918 µM O2 d−1
r2 = 0.838
Aug 26 : 5 m unfiltered
−1.275 µM O2 d−1
r2 = 0.692
Aug 26 : 5 m unfiltered
−2.254 µM O2 d−1
r2 = 0.833
Concentration (µM O2)
Incubation Time (hr)

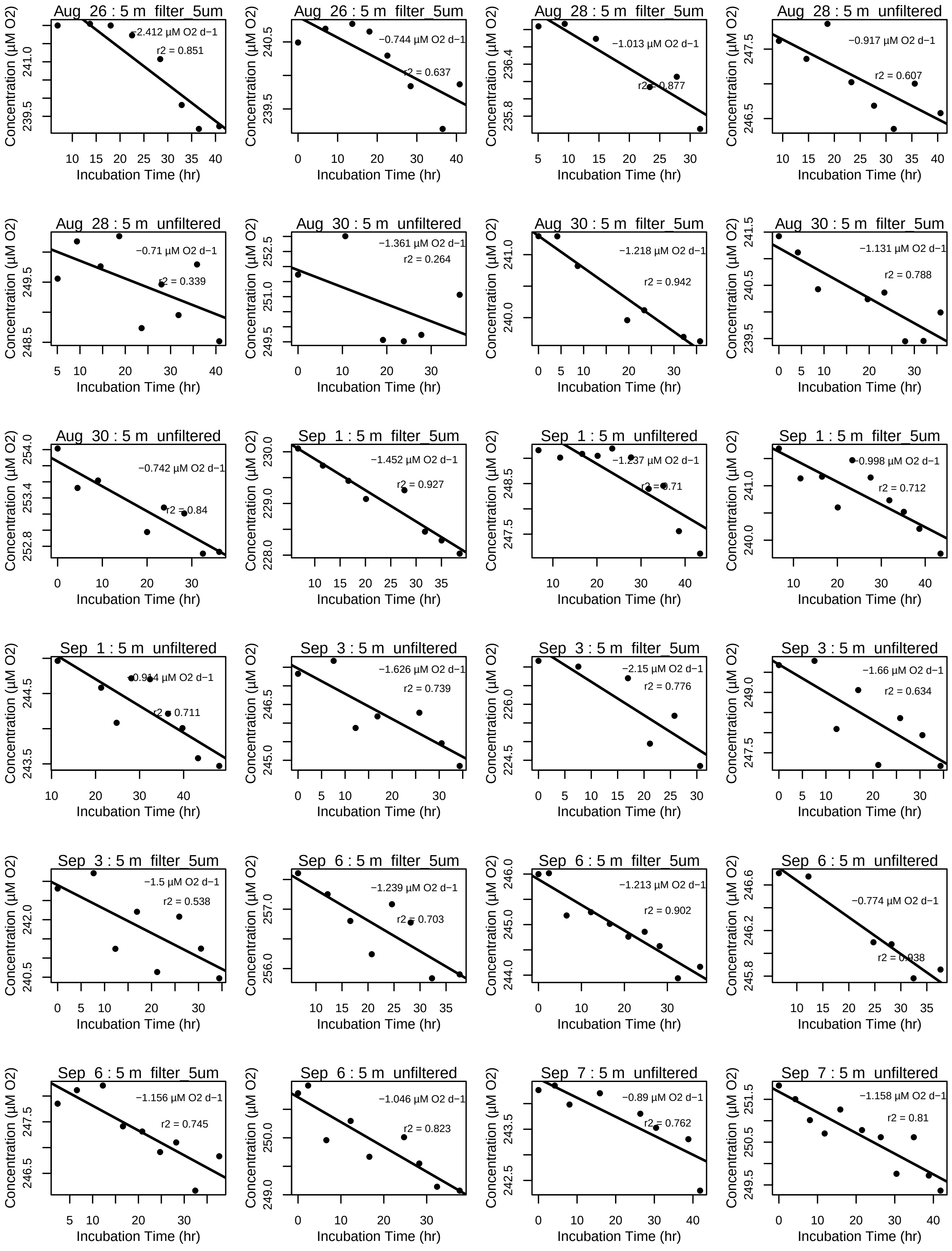
Aug 26 : 5 m filter_5um
−2.412 µM O2 d−1
r2 = 0.851
Aug 26 : 5 m filter_5um
−0.744 µM O2 d−1
r2 = 0.637
Aug 28 : 5 m filter_5um
−1.013 µM O2 d−1
r2 = 0.877
Aug 28 : 5 m unfiltered
−0.917 µM O2 d−1
r2 = 0.607
Aug 28 : 5 m unfiltered
−0.71 µM O2 d−1
r2 = 0.339
Aug 30 : 5 m unfiltered
−1.361 µM O2 d−1
r2 = 0.264
Aug 30 : 5 m filter_5um
−1.218 µM O2 d−1
r2 = 0.942
Aug 30 : 5 m filter_5um
−1.131 µM O2 d−1
r2 = 0.788
Aug 30 : 5 m unfiltered
−0.742 µM O2 d−1
r2 = 0.84
Sep 1 : 5 m filter_5um
−1.452 µM O2 d−1
r2 = 0.927
Sep 1 : 5 m unfiltered
−1.237 µM O2 d−1
r2 = 0.71
Sep 1 : 5 m filter_5um
−0.998 µM O2 d−1
r2 = 0.712
Sep 1 : 5 m unfiltered
−0.914 µM O2 d−1
r2 = 0.711
Sep 3 : 5 m unfiltered
−1.626 µM O2 d−1
r2 = 0.739
Sep 3 : 5 m filter_5um
−2.15 µM O2 d−1
r2 = 0.776
Sep 3 : 5 m unfiltered
−1.66 µM O2 d−1
r2 = 0.634
Sep 3 : 5 m filter_5um
−1.5 µM O2 d−1
r2 = 0.538
Sep 6 : 5 m filter_5um
−1.239 µM O2 d−1
r2 = 0.703
Sep 6 : 5 m filter_5um
−1.213 µM O2 d−1
r2 = 0.902
Sep 6 : 5 m unfiltered
−0.774 µM O2 d−1
r2 = 0.938
Sep 6 : 5 m filter_5um
−1.156 µM O2 d−1
r2 = 0.745
Sep 6 : 5 m unfiltered
−1.046 µM O2 d−1
r2 = 0.823
Sep 7 : 5 m unfiltered
−0.89 µM O2 d−1
r2 = 0.762
Sep 7 : 5 m unfiltered
−1.158 µM O2 d−1
r2 = 0.81
Concentration (µM O2)
Incubation Time (hr)

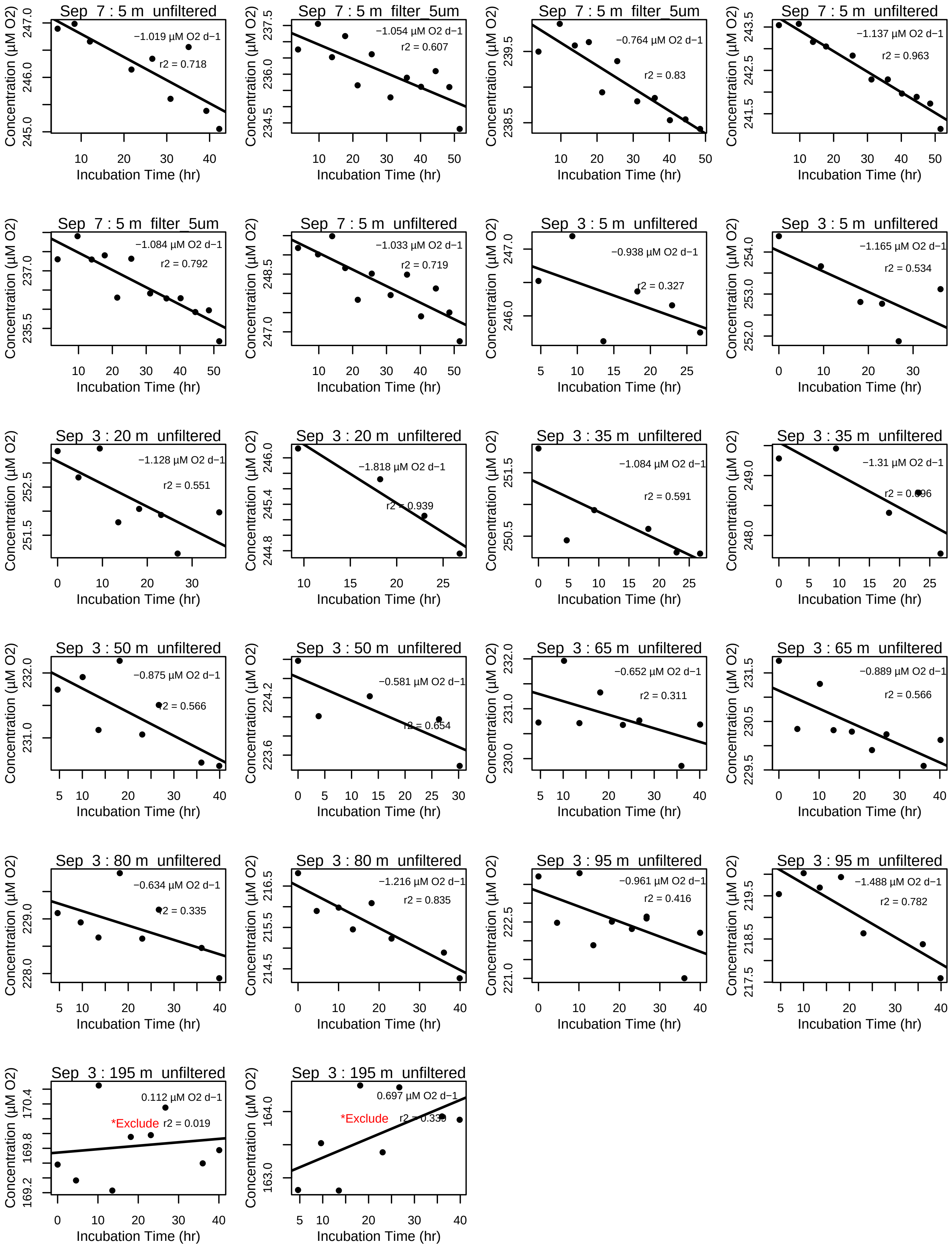
Concentration (µM O2)
Incubation Time (hr)
Sep 7 : 5 m unfiltered
−1.019 µM O2 d−1
r2 = 0.718
Sep 7 : 5 m filter_5um
−1.054 µM O2 d−1
r2 = 0.607
Sep 7 : 5 m filter_5um
−0.764 µM O2 d−1
r2 = 0.83
Sep 7 : 5 m unfiltered
−1.137 µM O2 d−1
r2 = 0.963
Sep 7 : 5 m filter_5um
−1.084 µM O2 d−1
r2 = 0.792
Sep 7 : 5 m unfiltered
−1.033 µM O2 d−1
r2 = 0.719
Sep 3 : 5 m unfiltered
−0.938 µM O2 d−1
r2 = 0.327
Sep 3 : 5 m unfiltered
−1.165 µM O2 d−1
r2 = 0.534
Sep 3 : 20 m unfiltered
−1.128 µM O2 d−1
r2 = 0.551
Sep 3 : 20 m unfiltered
−1.818 µM O2 d−1
r2 = 0.939
Sep 3 : 35 m unfiltered
−1.084 µM O2 d−1
r2 = 0.591
Sep 3 : 35 m unfiltered
−1.31 µM O2 d−1
r2 = 0.696
Sep 3 : 50 m unfiltered
−0.875 µM O2 d−1
r2 = 0.566
Sep 3 : 50 m unfiltered
−0.581 µM O2 d−1
r2 = 0.654
Sep 3 : 65 m unfiltered
−0.652 µM O2 d−1
r2 = 0.311
Sep 3 : 65 m unfiltered
−0.889 µM O2 d−1
r2 = 0.566
Sep 3 : 80 m unfiltered
−0.634 µM O2 d−1
r2 = 0.335
Sep 3 : 80 m unfiltered
−1.216 µM O2 d−1
r2 = 0.835
Sep 3 : 95 m unfiltered
−0.961 µM O2 d−1
r2 = 0.416
Sep 3 : 95 m unfiltered
−1.488 µM O2 d−1
r2 = 0.782
Sep 3 : 195 m unfiltered
0.112 µM O2 d−1
*Exclude
r2 = 0.019
Sep 3 : 195 m unfiltered
0.697 µM O2 d−1
*Exclude
r2 = 0.339

**Supplemental Figure 1.** Individual N. Pacific respiration assays grouped by sampling day, depth, and size fraction from the using PreSens oxygen sensor spots manually measured every few hours. The respiration rate is the linear regression (black line) of oxygen concentrations (black points). Assays with rates below detection or with nonlinear rates are indicated on the plot with the label "Excluded".

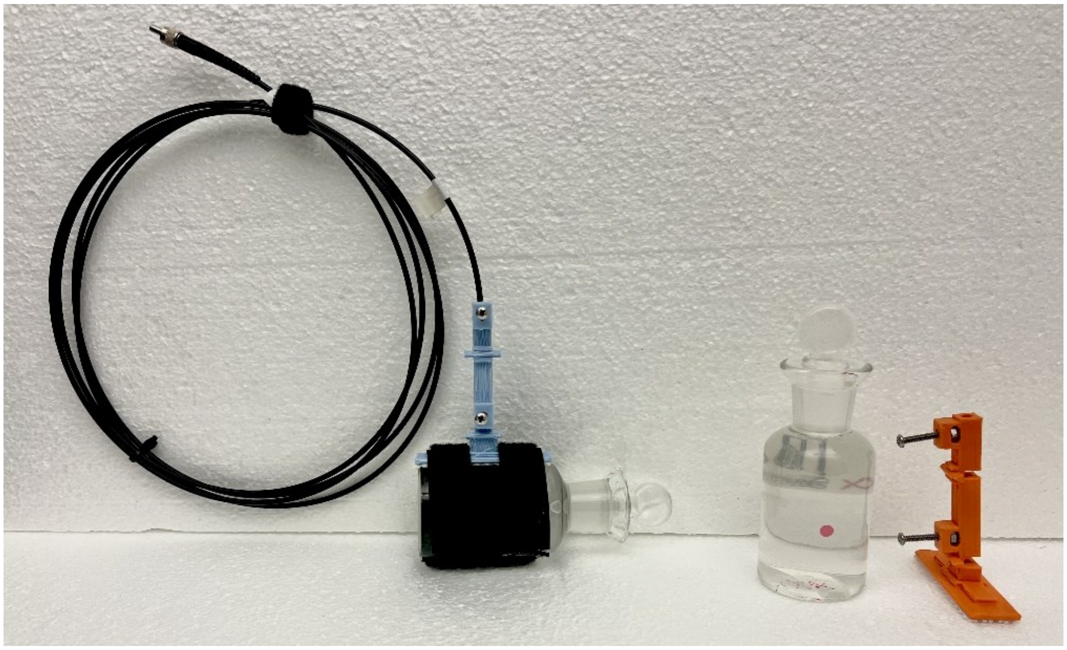

**Supplemental Figure 2.** Respiration incubation bottle setup from the N. Atlantic assays. Biological oxygen demand (BOD) bottles are filled with no headspace with sample water. A mount holds the polymer optical fiber (POF) to the exterior of the BOD wall such that the POF is aligned with the PreSens PSt3 foil sensor spot (pink) inside the BOD bottle. The holder has set screws which are gently hand-tightened to secure the POF. The base of the holder has rubber grip to prevent slipping when secured to the BOD bottle with a Velcro wrap. BOD bottles are placed in an opaque, temperature-controlled water bath with the sensor spot facing upwards as on the left. The bath is sealed and covered with a black bag with the POFs fed through a small hole and run to the controller boxes and computer. (POF Holder design available at https://github.com/2mrcohn/pub).

### North Atlantic Oxygen Drawdown Assays: May 06 , 2021

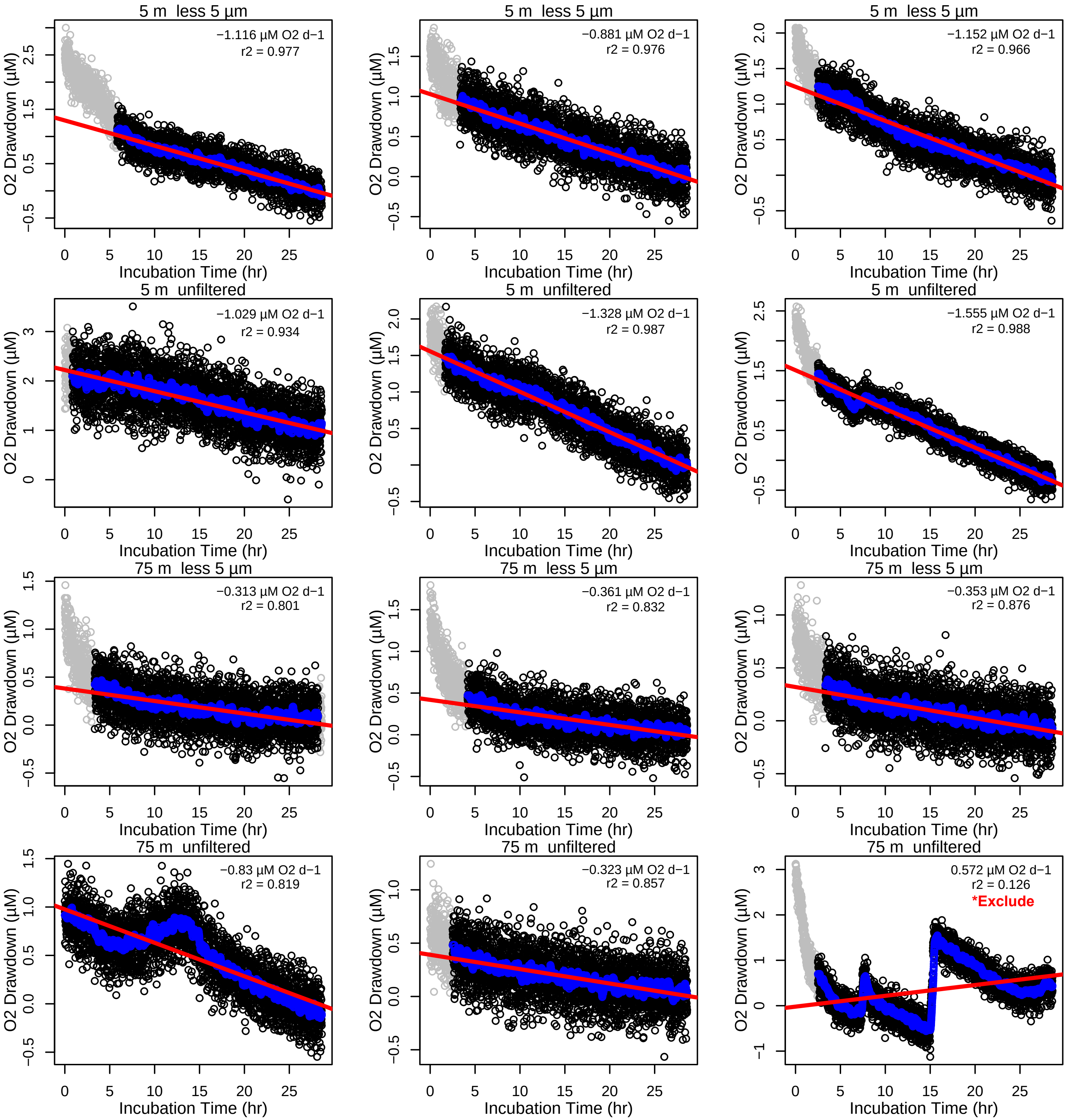

### North Atlantic Oxygen Drawdown Assays: May 12 , 2021

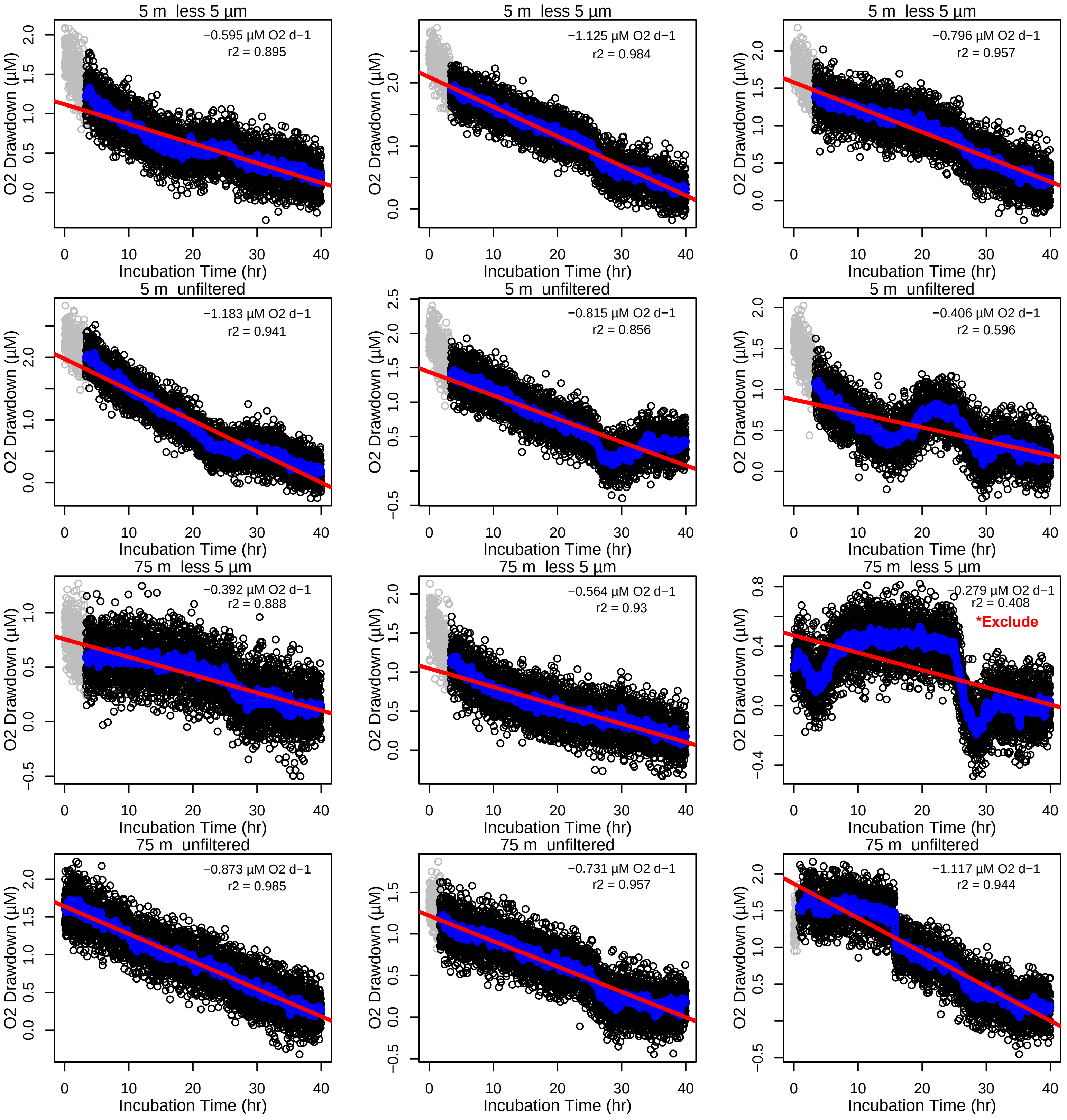

### North Atlantic Oxygen Drawdown Assays: May 14 , 2021

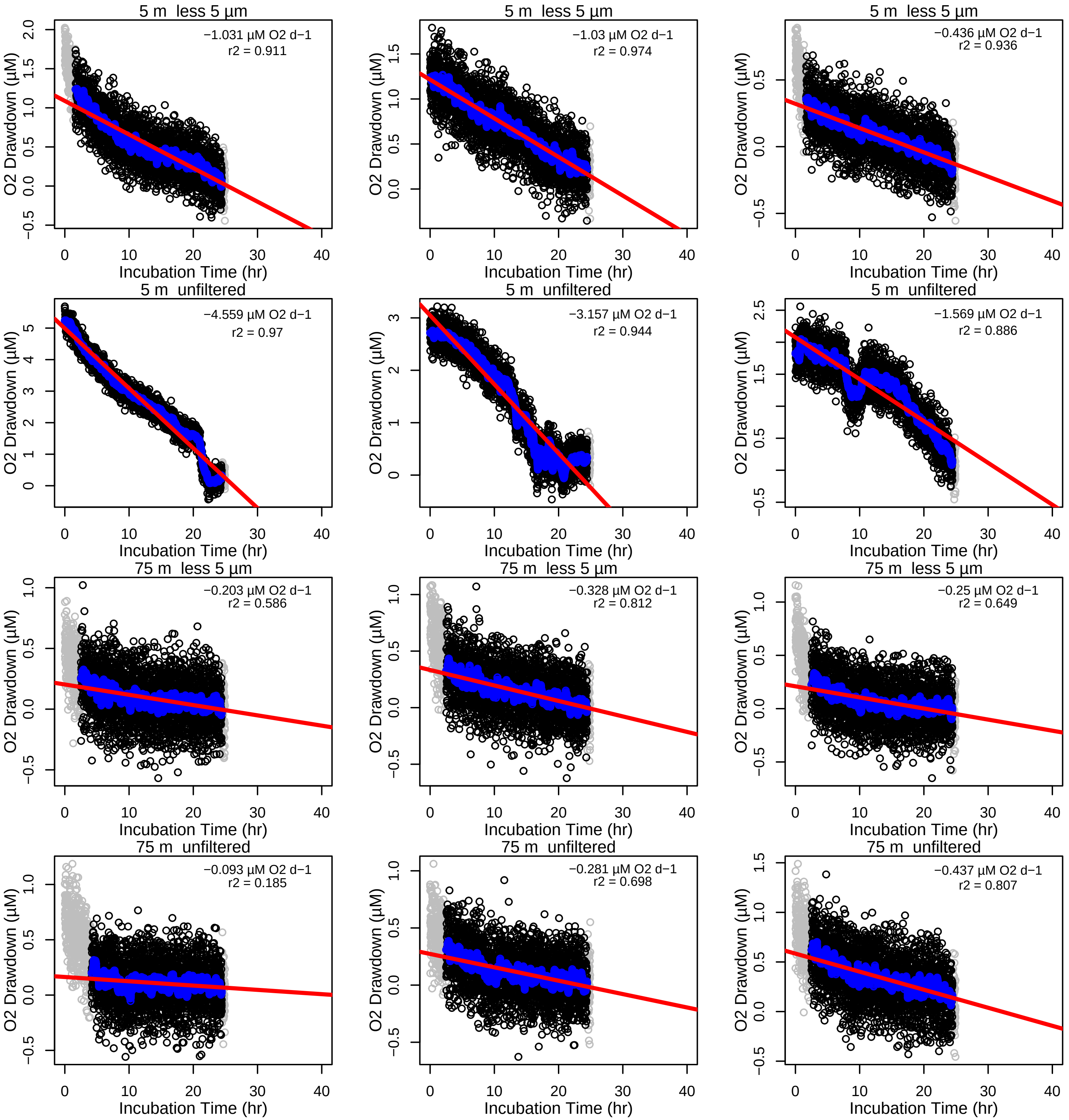

### North Atlantic Oxygen Drawdown Assays: May 18 , 2021

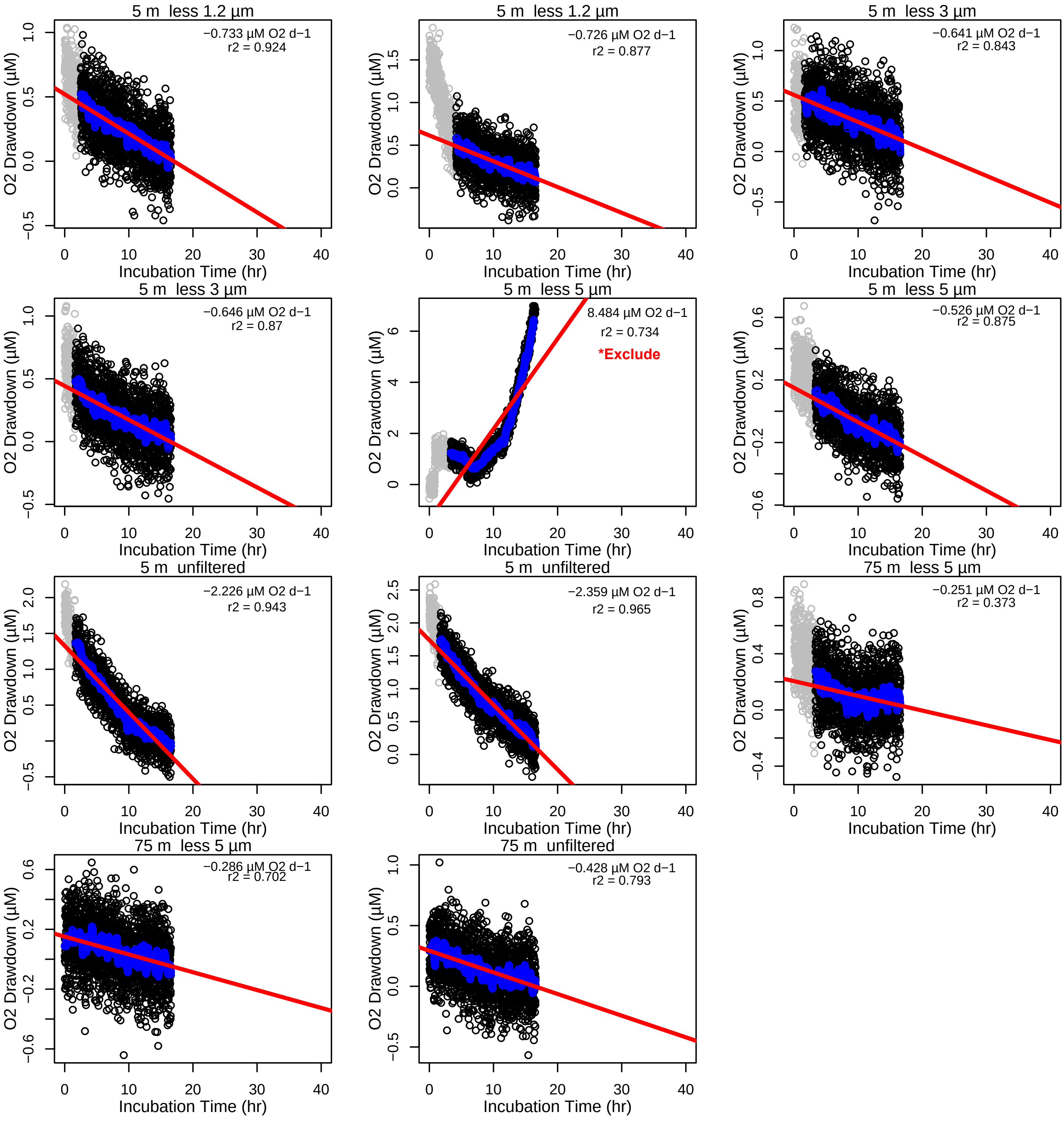

### North Atlantic Oxygen Drawdown Assays: May 19 , 2021

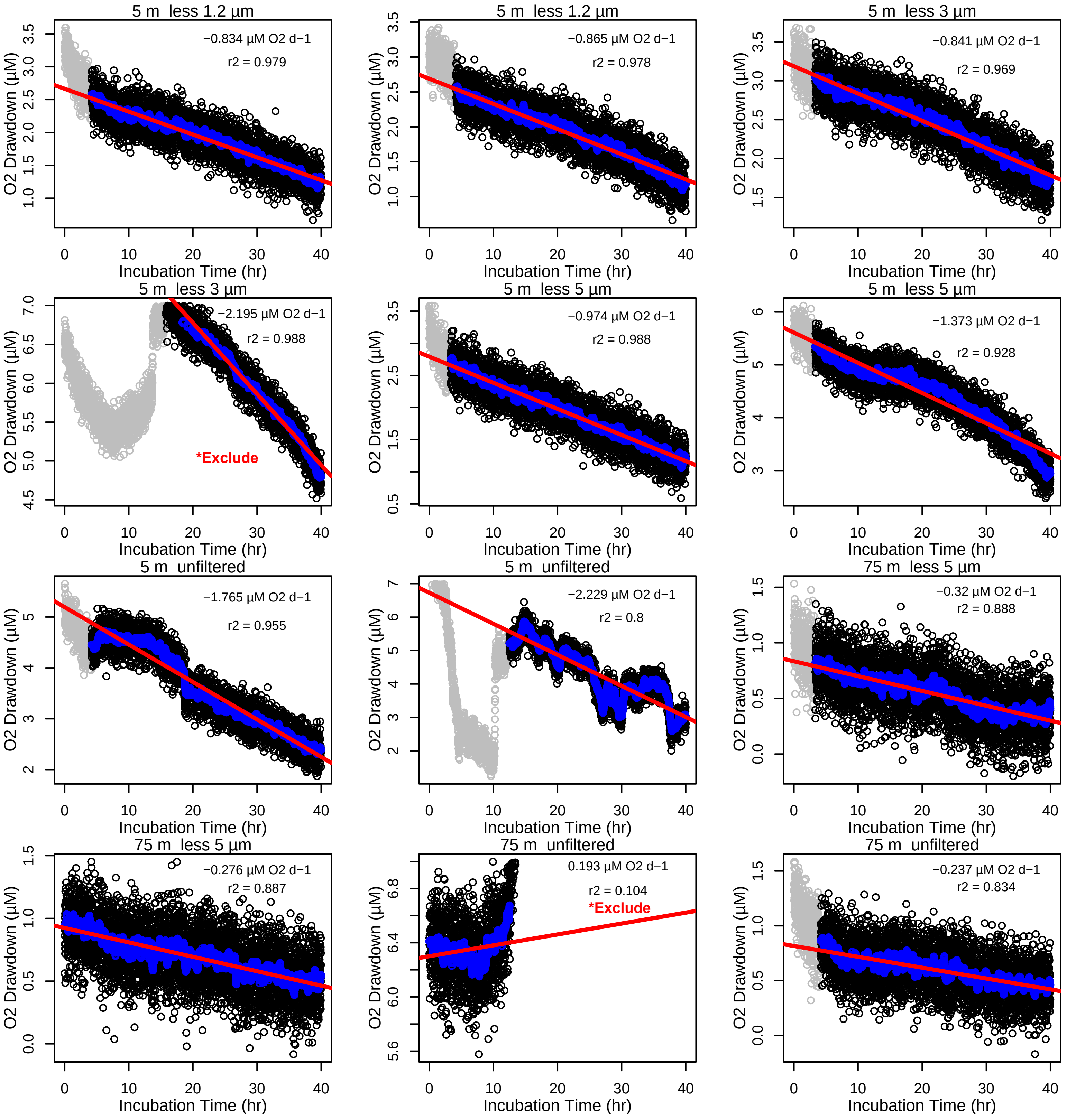

### North Atlantic Oxygen Drawdown Assays: May 22 , 2021

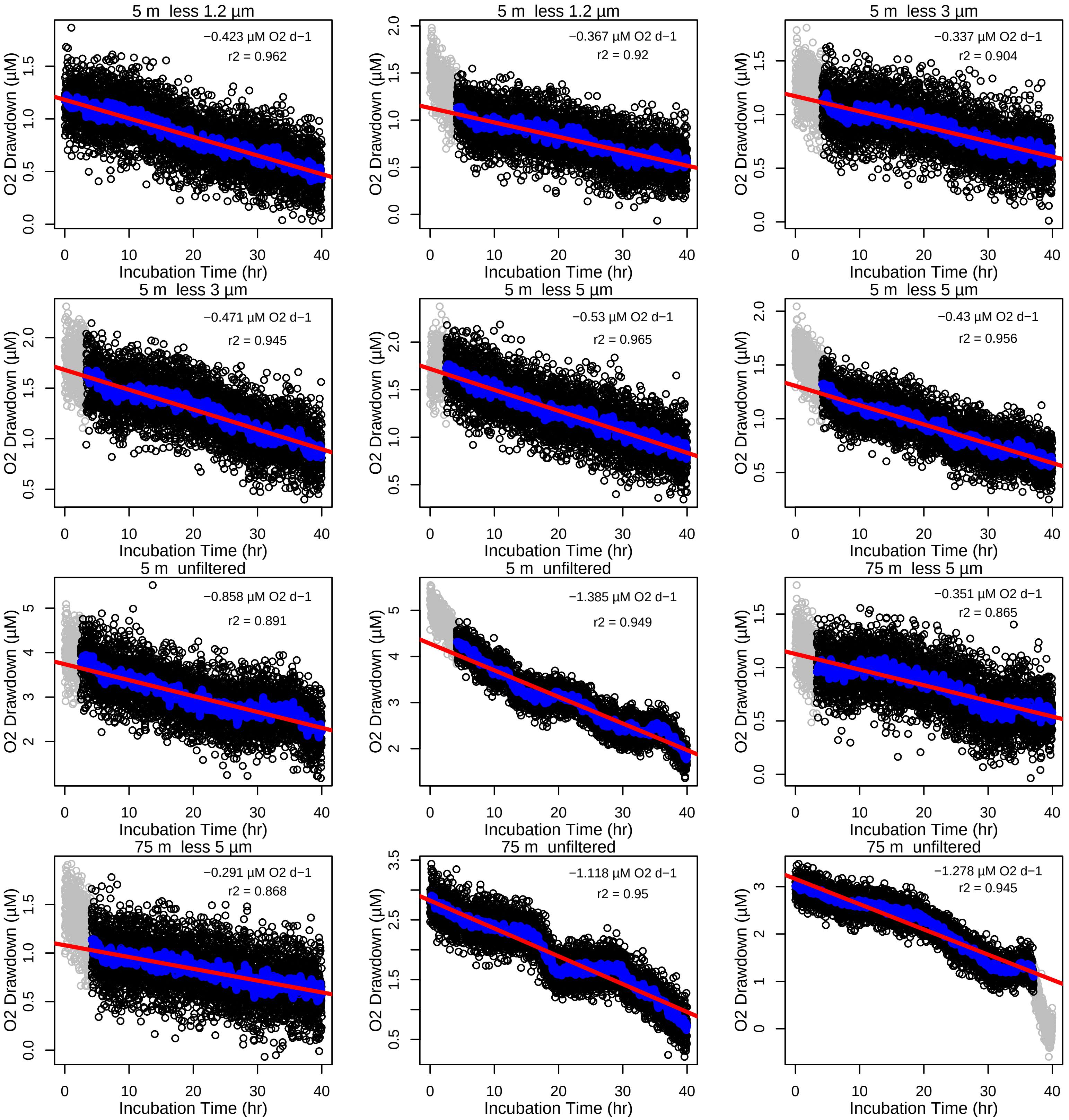

### North Atlantic Oxygen Drawdown Assays: May 25 , 2021

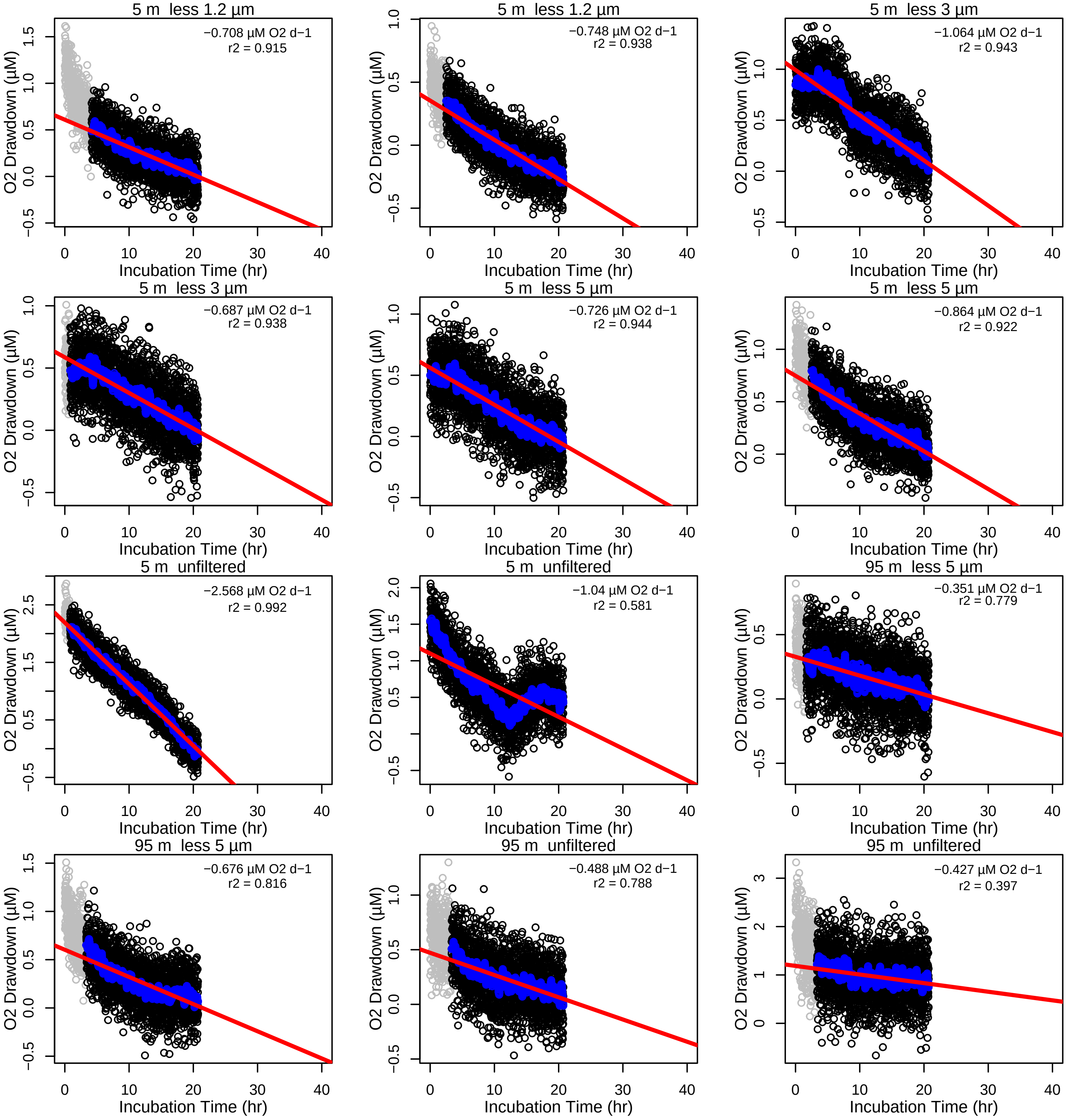

### North Atlantic Oxygen Drawdown Assays: May 26 , 2021

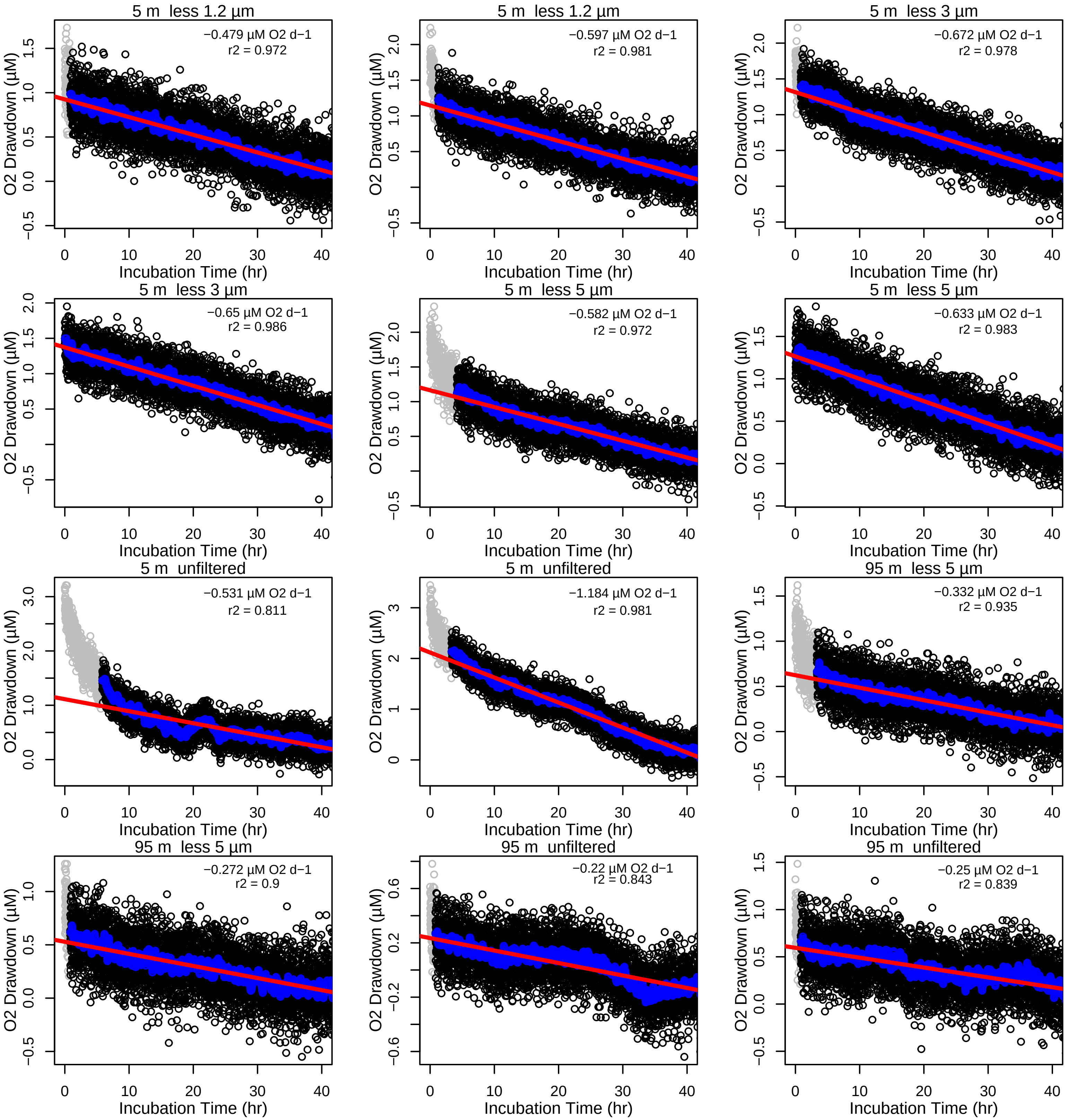

### North Atlantic Oxygen Drawdown Assays: May 28 , 2021

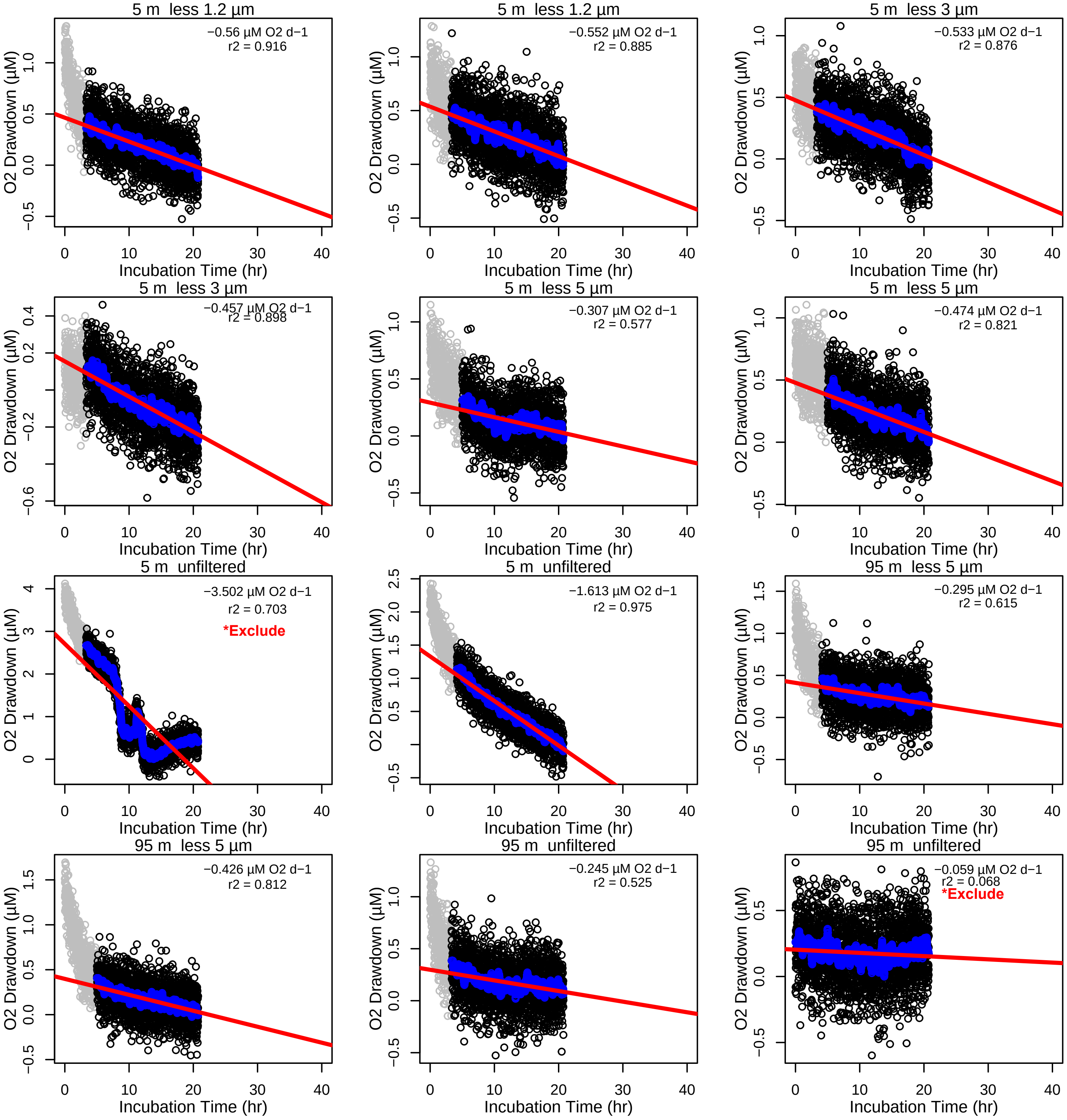

5m less 1.2 µm, Incubation
North Atlantic Oxygen Drawdown Assays: Diel Response to Phyto−DOM

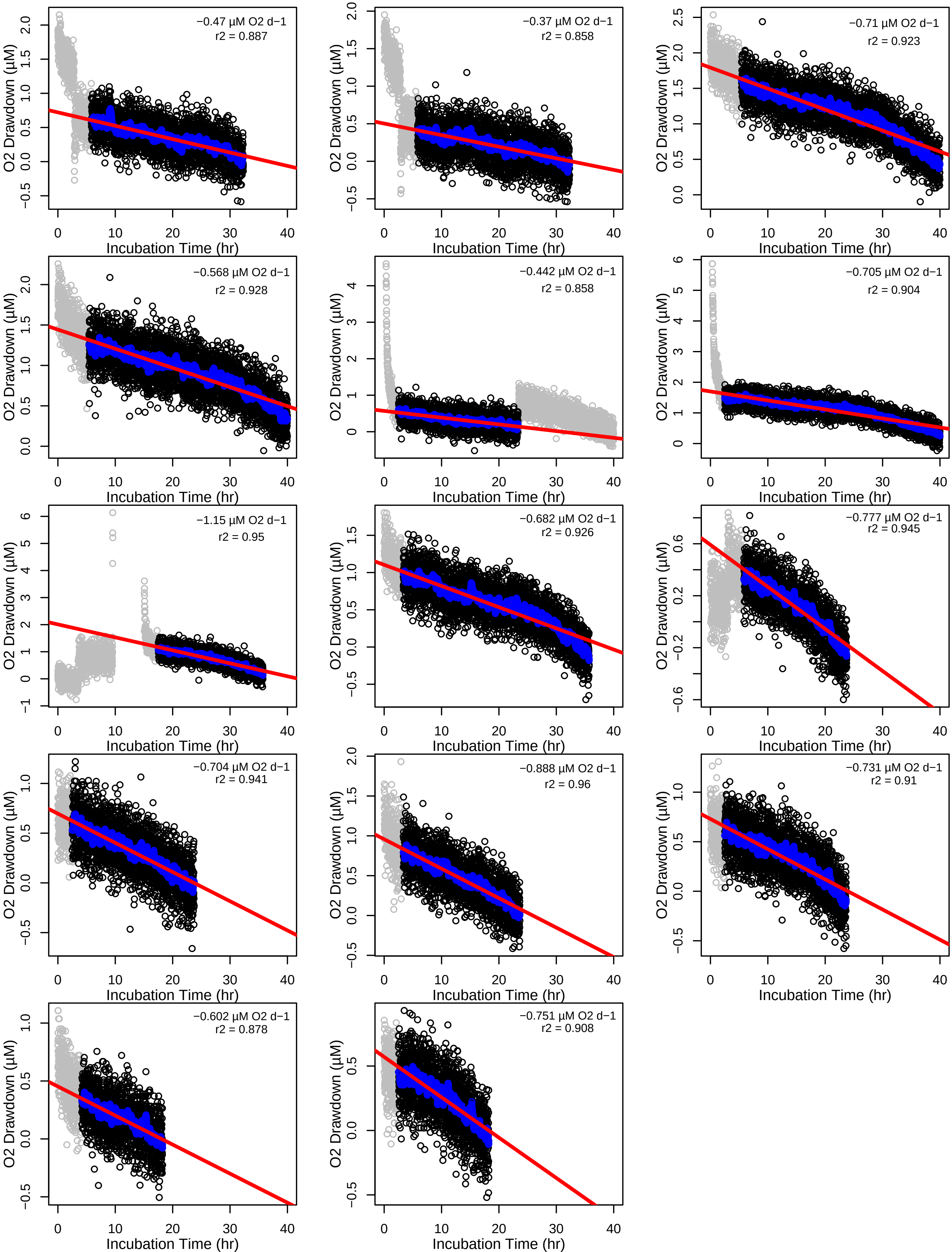

### North Atlantic Oxygen Drawdown Assays: O2 to INT Comparisson

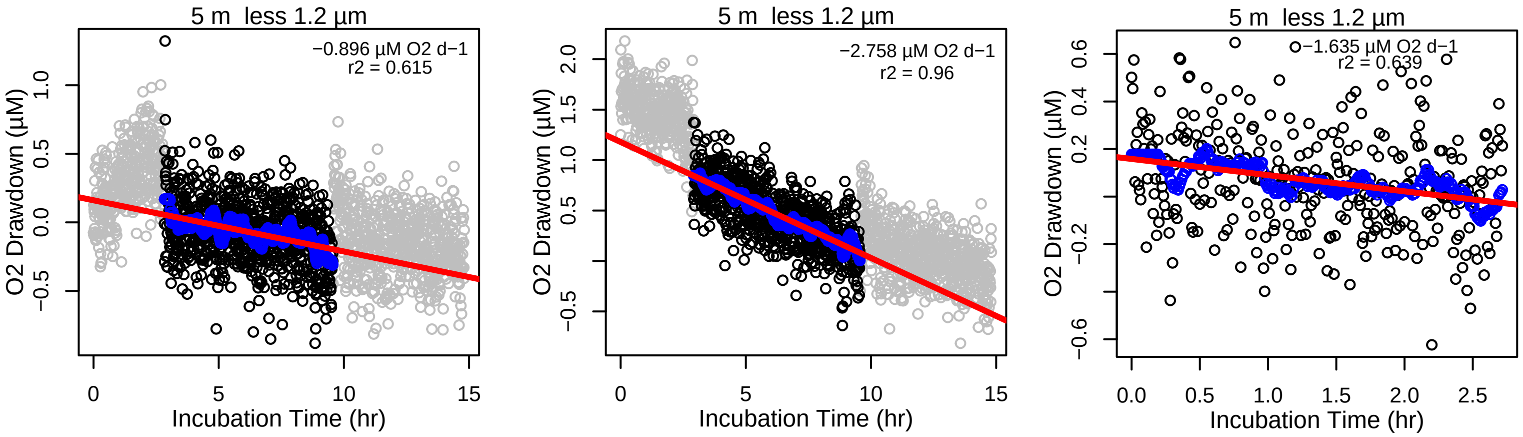

**Supplemental Figure 3.** Individual N. Atlantic respiration assays grouped by sampling day, depth, and size fraction using continuously mounted fiber optic cables to PreSens oxygen sensor spots. Oxygen concentrations were recorded every 30 s (black points) with removed measurements indicated (gray points). The rolling mean (blue points) and linear regression (red line, statistics in top right corners) show the respiration rate. Assays with rates below detection or with nonlinear rates are indicated on the plot with the label "Excluded".

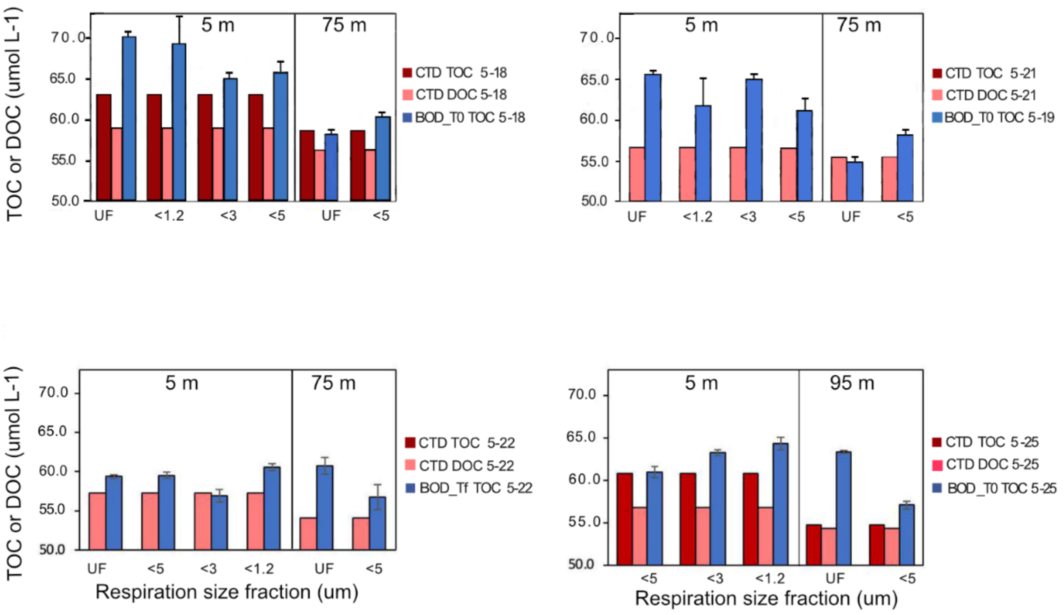


**Supplemental Figure 4.** Total (TOC) and dissolved organic carbon (DOC) values of select respiration incubations compared to CTD casts. Error bars indicate replicate values in the top row and instrumental error in the bottom row. One notable challenge when conducting microbial assays in oligotrophic environments is DOC contamination. In our N. Atlantic assays, a small (2 to 3 $\mu$mol $L^{-1}$) yet consistent increase in organic carbon was observed in the respiration assays' onset compared to Niskin seawater. However, since most of our measured rates were <1 $\mu$mol $O_2$ $L^{-1}$ $d^{-1}$ and we could clearly differentiate rates in the surface, sub-euphotic zones, and across various size classes, there was minimal evidence to suggest that the potential increase in organic carbon was labile. Further research is required to more accurately quantify the extent of DOC contamination during the bottle setup process, identify potential sources, and devise strategies to mitigate it.

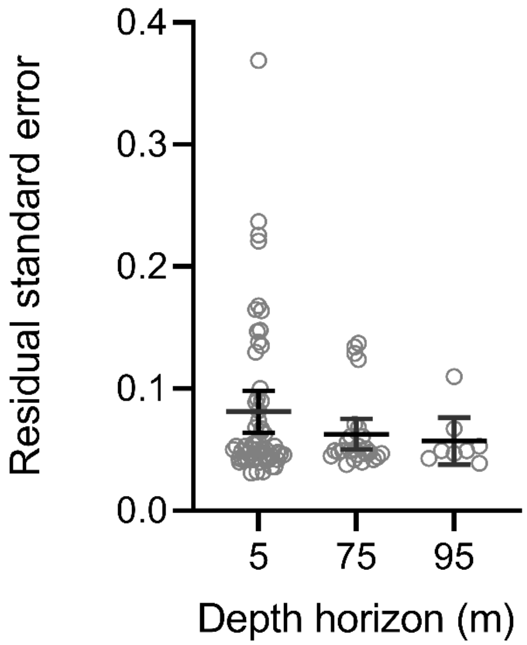


**Supplemental Figure 5.** Supplemental Figure 5. Residual standard errors (RSE) of the trimmed N. Atlantic respiration rates. Gray circles indicate the residual standard error for each N. Atlantic assay. The black bars show the mean and 95% confidence interval of the RSEs for each depth treatment. There is no significant difference between the RSE based on depth.

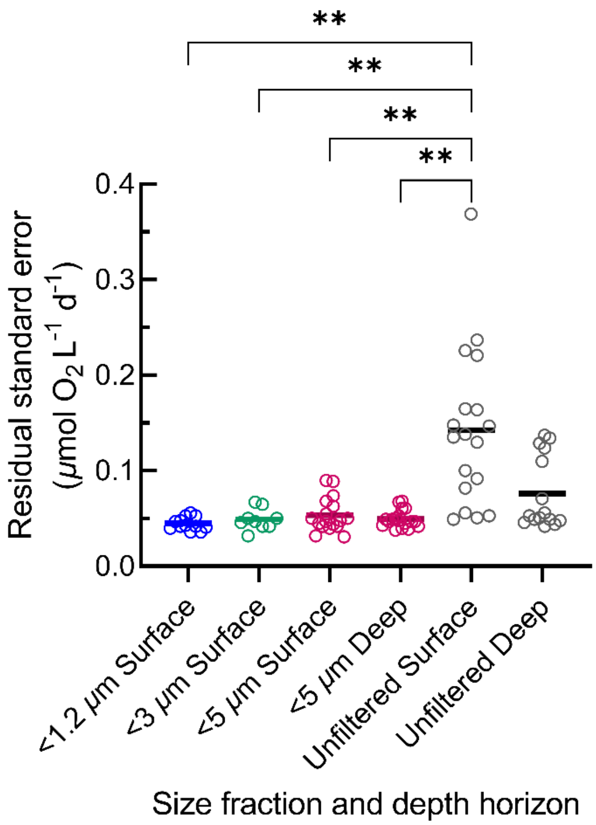


**Supplemental Figure 6.** Residual Standard Error (N. Atlantic) by size fraction. Horizontal bars indicate the mean RSE within each size class and depth horizon. One outlier was identified in the <5 $\mu$m surface fraction using the ROUT method (Q=1%) and removed. No outliers were identified for the <1.2 $\mu$m fraction (n=12), the <3 $\mu$m fraction (n=9), the <5 $\mu$m deep fraction (n=18), or the unfiltered surface and deep fractions (n=18 and n=16). Means of each fraction were compared using a Brown-Forsythe or Welch ANOVA with significant differences among means at $p<0.001$ indicated (**).

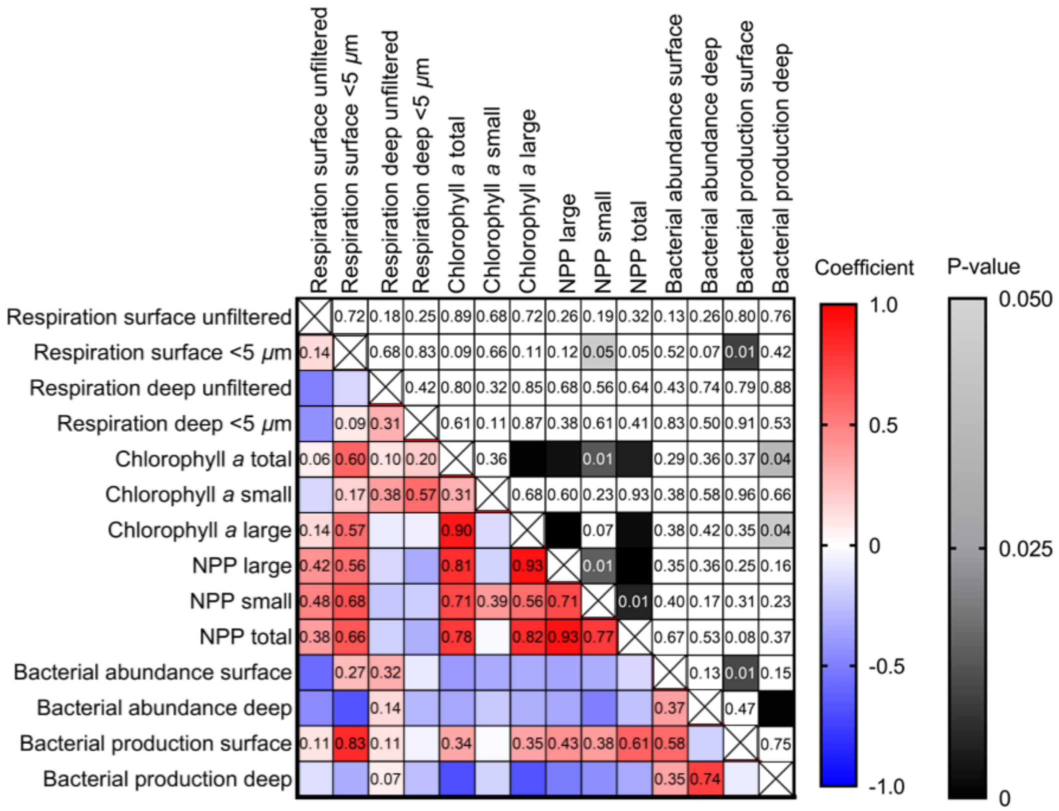


**Supplemental Figure 7.** Matrix of Pearson's correlation coefficients (bottom diagonal) and p-values (top diagonal) to identify statistically significant relationships ($p<0.05$, gray shaded cells) between biological variables in the N. Atlantic.

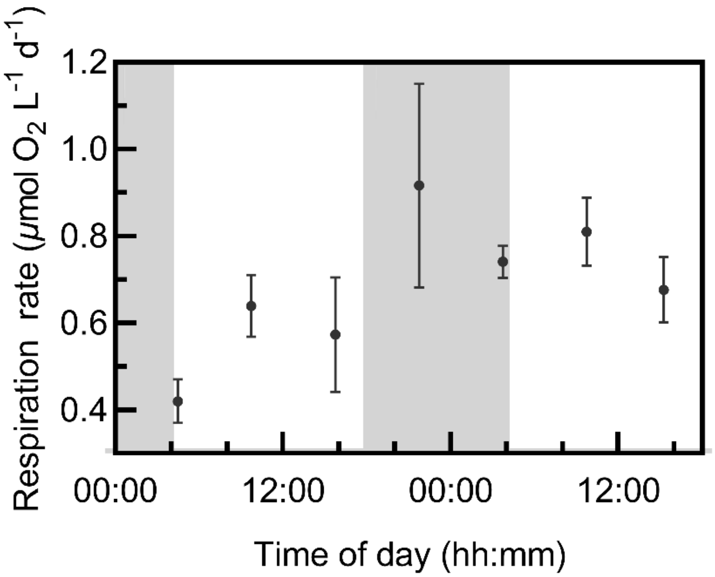


**Supplemental Figure 8.** Mean (n=2) and range of <1.2 $\mu$m respiration rate from on-deck incubations targeting the response of bacterial respiration to phytoplankton-derived dissolved organic matter over a diel period. Night periods (dusk to dawn) are shaded in gray.
